# Genome-scale perturbation signatures from primary human CD4^+^ T cells improve genetics-based prioritization of immune drug targets

**DOI:** 10.64898/2026.09.12.748195

**Authors:** Alexander L. Han, Max Slotnik, David A. Fox, Ryan S. Dhindsa, Johann E. Gudjonsson, J Michelle Kahlenberg, Joshua Welch

## Abstract

While human genetic evidence improves drug program success, target selection largely relies on associational features. Even functional features are mainly observational, capturing disease associations rather than the consequence of perturbing genes in the disease-relevant human cell type. To move beyond association, we present IGNITE (Immune Genomics and fuNctional Integration for Target Enrichment), a framework that prioritizes immune drug targets by integrating human genetic priors with features from genome-scale perturb-seq in 22 million primary human CD4^+^ T cells and polarized T-helper subset differential expression. IGNITE is a semi-supervised machine learning model that applies positive-unlabeled learning to approved immune targets, then ranks 19,502 protein-coding genes to prioritize new candidates. In 727 in-trial genes held out from training, functional genomics increased target enrichment among the top 50 nominations from 2.7- to 4.8-fold for immune trial targets at any phase, and from 4.5- to 5.9-fold for targets in Phase III. These performance gains were immune-specific, as IGNITE outperformed genetic comparators on predicting immune-exclusive targets but not on cardiac-exclusive targets. Using temporal validation with labels frozen in 2014, IGNITE outperformed the genetics-only model in ranking 134 genes that subsequently entered immune trials. The functional genomics layer also surfaced two novel, pharmacologically tractable candidates, *ELOVL6* and *RUVBL1*, elevating their rankings from outside the top 1,000 into the top 50. These findings highlight that perturbational signatures in the disease-relevant human cell type harbor target-relevant signals beyond human genetics, offering a generalizable route to indication-specific prioritization as perturbation atlases expand. Full IGNITE scores are publicly available at https://ignite.eecs.umich.edu/

## Introduction

Human genetic evidence has become one of the most valuable predictors of drug target success, but its predictive value is not uniform. Targets with genetically supported mechanisms are substantially more likely to reach approval^1^. Human genetic support was present in roughly two-thirds of drugs approved by the FDA in 2021, and genetic associations can also predict safety liabilities that lead to clinical trial discontinuations^2,3^. The probability of drug program success is associated with confidence in the causal gene, which is greatest for Mendelian traits, rare coding variants, and GWAS associations linked to coding variants^4–7^. The predictive signal in human genetics is thus driven primarily by resolution of the causal gene rather than by association strength. For most genes, available evidence is limited to association studies that link a trait to a locus rather than a gene, do not consider the cell type or state, and involve common variants with smaller effect sizes where the causal gene is especially difficult to identify.

Scores such as Genetic Priority Score (GPS) and ML-GPS inherit this limitation because they are built almost entirely on statistical genetic evidence^8,9^. Others including Priority Index (PI) and Polygenic Priority Score (PoPS) incorporate curated annotations such as ontology terms and network connectivity^10–12^, which are vulnerable to coverage bias because annotation depth tracks how extensively a gene has already been studied^13–15^. Since well-studied genes are also enriched among approved targets used to train and benchmark these models, this same bias can inflate performance estimates^16–18^. These approaches differ in architecture but share a constraint. All input features lack the experimental interpretation of downstream consequences of perturbing a specific gene in disease-relevant human cells^19,20^. Closing this gap requires focusing on a disease domain where the major causal cell types are defined and where genome-scale perturbation data for the disease-relevant cell type are available. Currently, immune-mediated inflammatory diseases (IMIDs) and CD4^+^ T cells meet these criteria.

IMIDs collectively affect roughly 5-7% of the population and caused 67.6 million incident cases globally in 2019^21,22^. Despite more than 30 approved targeted immunomodulators, many patients with IMIDs remain refractory to existing therapies. For instance, 30-40% of patients with rheumatoid arthritis or spondyloarthritis cannot achieve durable clinical benefit, and anti-TNF primary non-response accounts for 20-40% of cases in Crohn’s disease^23–26^. Thus, there is clear clinical demand for mechanistically distinct targets.

IMID indications also offer an unusual advantage for testing whether perturbation signatures add complementary signals to established genetic priors. The causal cell type is largely specified, with CD4^+^ T cells disproportionately enriched for immune-mediated disease variants relative to other cell types^27–30^. In addition, perturbation of regulatory genes and CRISPR screens in primary human T cells reveal immune network architecture and suggest that responses in these immune cells are stimulation-dependent^19,31^. These perturbational functional genomics efforts have extended to genome-scale perturb-seq in primary human CD4^+^ T cells at rest and post-stimulation^32^. We distinguish two complementary forms of functional genomics: observational, which captures differential expression across polarized T-helper states without perturbation, and perturbational, which captures transcriptional changes from directly perturbing each gene through perturb-seq.

With perturbation phenotypes of CD4^+^ T cells available at genome scale, it is possible to explore whether perturbational functional genomics and genetic priors can be integrated effectively. Recent work at this intersection includes single-cell CRISPR screens that map target genes at GWAS loci, and a machine-learning model developed to identify targets restricted to immuno-oncology^33,34^. Latest efforts have integrated genetic evidence with perturbation data to infer disease mechanisms^35^. However, to our knowledge, no supervised or semi-supervised model has been trained to prioritize therapeutic targets by combining human genetic priors with genome-scale experimental perturbation responses from the relevant primary human cell type.

Existing immune-focused prioritization methods maximize recall of known targets by aggregating genetic, regulatory, and extensive literature-derived annotation evidence^10,11^. Here, we instead test whether observational and perturbational functional genomics improve prioritization of immune targets beyond human genetic evidence, a distinct question that a target recall benchmark is not built to measure. The natural approach is positive-unlabeled (PU) learning because only approved targets are reliable positives while unlabeled genes cannot be assumed to be negatives^36,37^. The resulting model, IGNITE (Immune Genomics and fuNctional Integration for Target Enrichment), spans 19,502 genes and combines genetic priors including association with inborn errors of immunity (IEI), genic intolerance estimates, and common and rare variant association evidence with CD4^+^ T-cell perturb-seq and polarized T-helper subset differential expression^32^.

## Results

### Construction of the genome-scale gene-evidence dataset

IGNITE prioritizes drug targets for immune-mediated disease across the protein-coding genome (**Figure 1**). The feature matrix spans 19,502 genes and carries 24 total features from three categories (**see Methods, Supplementary Table 2**). The first category, genetic features, includes: IEI gene membership^38^ (Mendelian), composite GWAS score and composite gene burden score for immune-mediated disease^39^ (common- and rare-variant association), and two genic intolerance estimates^40^ (disease-agnostic constraint). Observational functional genomics, the second category, includes four polarized T-helper subset differential-expression z-scores where Th1, Th2, Th17, and Treg cells are each compared against resting Th0 cells^32^. The third category, perturbational functional genomics, comprises 15 features derived from genome-scale perturb-seq in 22 million primary human CD4^+^ T cells^32^.

**Figure 1.**
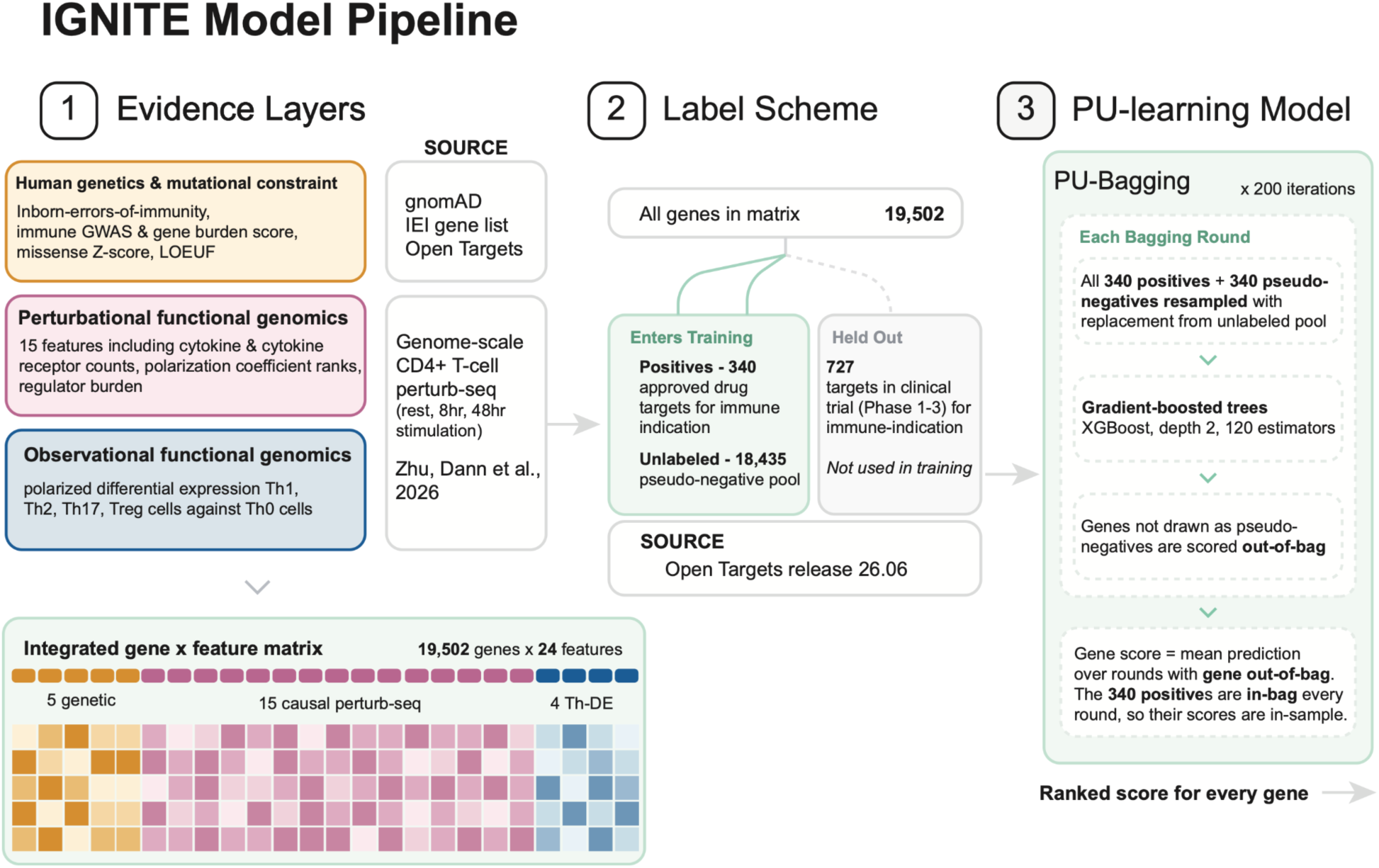
Overview of IGNITE framework for prioritizing immune drug targets. Schematic of IGNITE pipeline, showing three evidence blocks, the immune-indication label scheme, and the PU bagging model used to score genes (see Methods).

Each gene was labeled by the most advanced trial stage reached by any immune-indication drug targeting it in Open Targets^39^: approved, in-trial (Phase I/II/III), or non-target. After quality control, this labeling yielded 340 approved immune targets and 734 in-trial genes, of which 727 are present in the feature matrix (**see Methods, Supplementary Table 3**). The 340 approved targets are training positives. The 727 in-matrix in-trial genes are held out as a validation set and do not enter the training phase. Prioritization scores in IGNITE are computed from positive-unlabeled bagging. Each bag draws pseudo-negatives exclusively from the unlabeled pool, which excludes all held-out in-trial genes (**see Methods**).

### Four independent genetic axes distinguish immune drug targets

The four distinct lines of genetic evidence encompass Mendelian immune diseases, common-variant GWAS signals, rare-variant gene burden analyses, and metrics of genic intolerance (**Supplementary Table 5**). Approved and in-trial immune targets were strongly enriched for human immune genetic evidence and genic intolerance relative to the genome background (**Figure 2**). Genes associated with IEI were enriched among target classes with an odds ratio of 9.0 (95% CI 6.6-12.2) for approved targets and 3.3 (2.4-4.4) for in-trial targets against the genome (**Figure 2a, Supplementary Table 4**). Against a druggable genome background^41^, the corresponding odds ratios were 5.7 (4.1-8.0) and 2.1 (1.5-2.9), respectively (**Figure 2a**).

**Figure 2.**
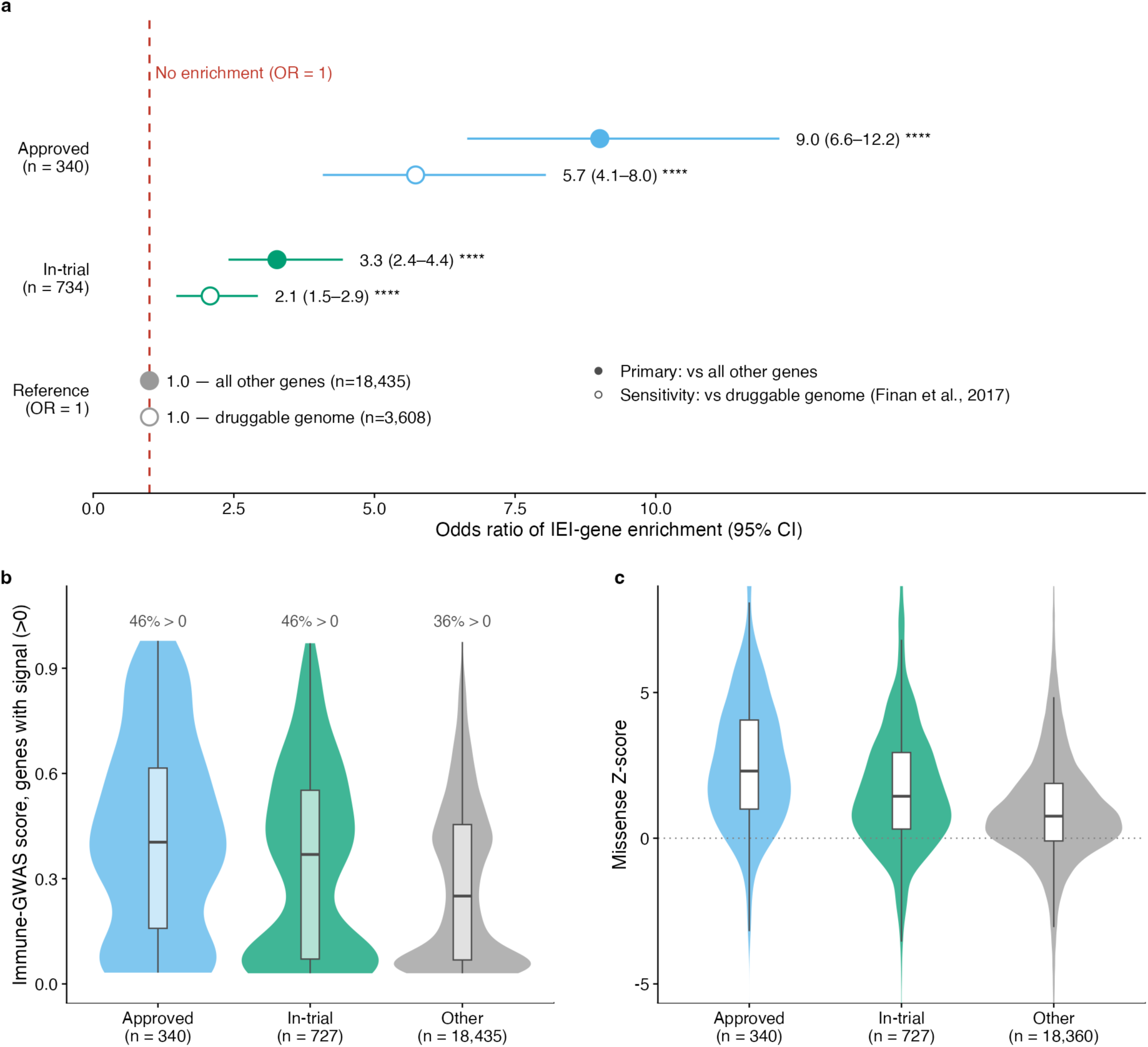
Genetic and intolerance evidence separates immune drug targets. **(a)** Odds ratio (±95% CI) for IEI-gene enrichment among approved and in-trial targets computed against all other genes and against the druggable genome as background. The dashed line indicates no enrichment. Group differences assessed by two-sided Fisher exact test. **(b)** Immune-GWAS score across approved, in-trial, and all other genes, restricted to genes with signal (>0). Percentages above indicate the fraction of genes with nonzero score. The n below reflects the full group size before this restriction. Violin shows distribution of scoring genes, and boxplot denotes median and interquartile range (IQR). In-trial targets are restricted to 727 genes after excluding 7 mitochondrially encoded genes. **(c)** gnomAD Missense Z-score compared across approved, in-trial, and all other genes. Violins with boxplot. 75 non-target genes lacked Missense Z-scores and were excluded from this panel, restricting the panel to 19,427 genes. * p < 0.05, ** p < 0.01, *** p < 10^−3^, **** p < 10^−4^; ns, not significant.

Approved and in-trial immune targets were also significantly more likely to harbor immune-GWAS signals, with 46% of approved and 46% of in-trial targets carrying a signal, compared to 36% for the remainder of the genome (**Figure 2b, Supplementary Table 4**). Among genes with immune-GWAS signal, the median immune-GWAS signal was 0.40 for approved targets, 0.37 for in-trial targets, and 0.25 for all other genes. In addition, targets exhibited greater intolerance to missense variants with a median Missense Z-score of 2.31 for approved targets, 1.44 for in-trial targets, and 0.76 for other genes (**Figure 2c**).

For both immune-GWAS signal and Missense Z-score, Kruskal-Wallis testing with pairwise Mann-Whitney U post hoc comparisons revealed that their monotonic increase from non-target to in-trial to approved genes was significant (**Supplementary Table 4**). In particular, IEI inclusion and Missense Z-score demonstrated a strong stepwise increase in enrichment from background to in-trial to approved genes. These findings support their use as genetic priors in the model alongside LOEUF, an estimate of loss-of-function (LOF) intolerance.

### Functional genomics in primary CD4^+^ T cells marks immune drug targets

Having established that genetic evidence separates immune drug targets, we next asked whether functional genomics in primary CD4^+^ T cells also carries target-relevant signals. We began with differential expression across polarized T-helper subsets. Th17 and Treg cells were the most transcriptionally distinct from resting Th0 with 48.1% and 55.5% of tested genes reaching 10% FDR relative to 21.4% in Th2 and 1.8% in Th1 (**Figure 3a, Supplementary Table 4**). Fisher’s exact test demonstrated that approved and in-trial targets were significantly more likely than non-target genes to achieve differential expression at 10% FDR in Th1 cells, though this pattern was not observed in Th2, Th17, or Treg cells (**Supplementary Table 4**). Trans-regulatory burden was also significantly elevated in approved and in-trial targets (**Figure 3b, Supplementary Table 4**). Kruskal-Wallis testing with pairwise Mann-Whitney U post hoc comparisons revealed a monotonic increase in residualized trans-regulatory burden from non-target to in-trial to approved genes. In contrast, cytokine and cytokine-receptor regulation did not show strong target-related enrichment (**Figure 3c**, **3d**). Across all three conditions, only a minority of genes significantly regulated cytokines or cytokine receptors, and this fraction was comparable between drug-status groups (**Supplementary Table 4**). The one exception was cytokine-receptor regulation at 8hr stimulation. Here, Fisher’s exact test showed that approved targets were significantly more likely than non-targets to regulate at least one receptor at 11.7% compared to 6.9% (**Supplementary Table 4**). Importantly, a gene can regulate cytokines or their receptors without carrying GWAS signals or Mendelian disease associations. Functional genomics therefore provides evidence for immune targets that is orthogonal to human genetic priors (**Supplementary Table 5**).

**Figure 3.**
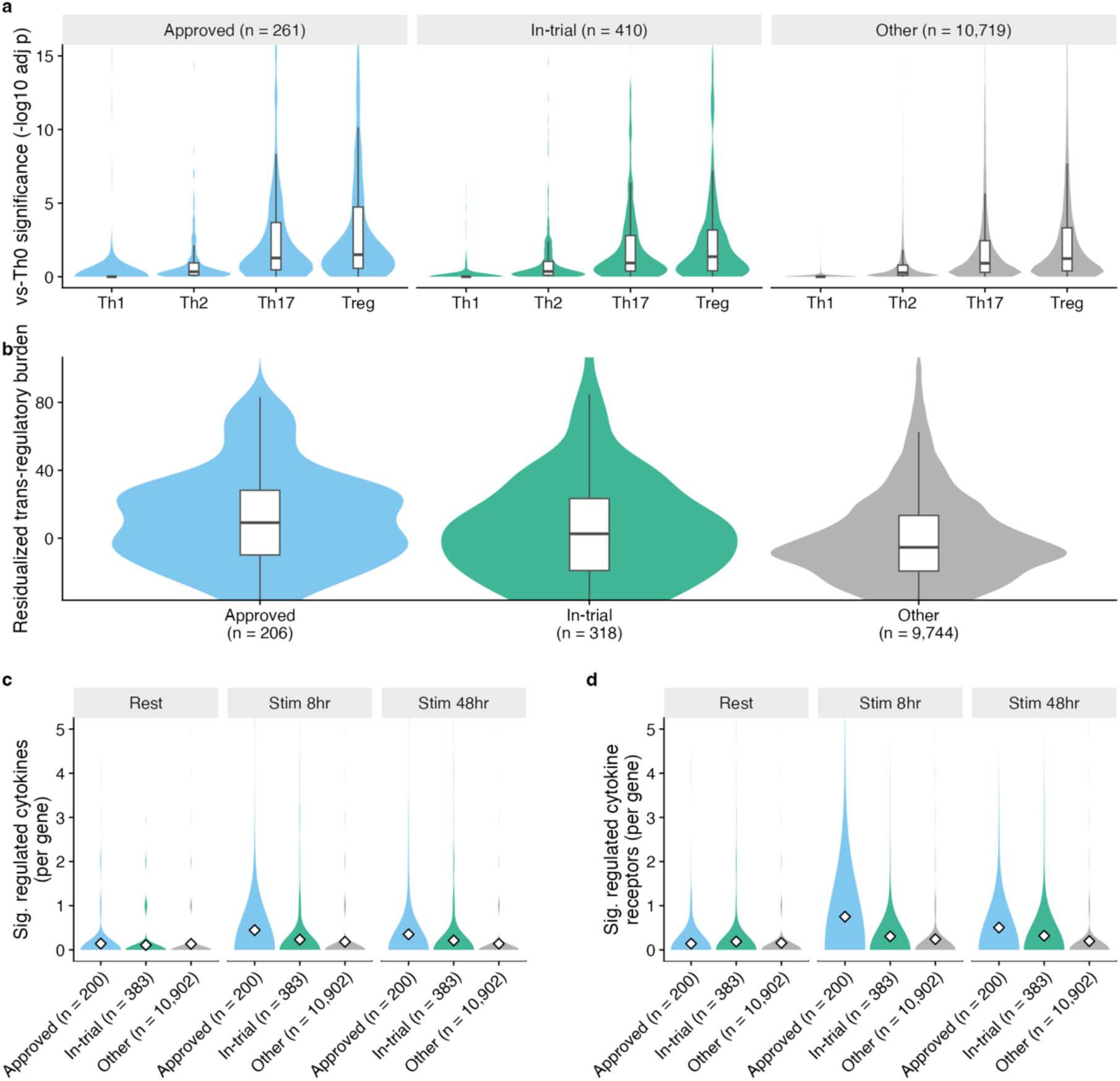
Observational and perturbational functional genomics in primary CD4^+^ T cells mark immune drug targets. **(a)** Differential expression significance of Th1, Th2, Th17, and Treg cells relative to Th0 across approved, in-trial, and all other genes. Values are computed from a per-gene Wald test with adjustment applied within each polarized subset. Genes without a measurable estimate in each subset are omitted. Boxplot denotes median and IQR. **(b)** Residualized trans-regulatory burden shown for approved, in-trial, and all other genes. Genes without a measurable estimate in each subset are omitted. **(c)** Number of significantly regulated cytokines per gene at rest, 8hr- and 48hr-stimulation, across approved, in-trial, and all other genes. Open diamonds denote group means. Genes without a measurable estimate in each subset are omitted. **(d)** Number of significantly regulated cytokine receptors per gene at rest, 8hr- and 48hr-stimulation, across approved, in-trial, and all other genes. Open diamonds denote group means. Genes without a measurable estimate in each subset are omitted.

### Functional genomics improves immune target prediction

To test whether these features improve prioritization beyond merely correlating with target status, we trained positive-unlabeled models on nested combinations of the three evidence blocks. Adding either functional genomics block outperformed the genetics-only model in predicting approved immune targets, with the full model achieving the highest cross-validated AUC (**Figure 4a, Supplementary Table 6**). The Nadeau–Bengio corrected t-test demonstrated that these gains against the genetics-only model were significant (**Figure 4c**). A grouped-Shapley decomposition of AUC-above-chance attributed 68% of the full model’s performance to the genetic block, 19% to the observational block, and 13% to the perturbational block (**Figure 4b**). The positive-unlabeled model score rose monotonically across target classes, with median cross-fitted rank percentile of 0.486 for unlabeled, 0.694 for held-out in-trial targets, and 0.875 for approved targets (**Figure 4d, Supplementary Table 7**). The 727 in-trial targets that were held out from model training ranked significantly above the rest of the unlabeled genes. The full model also recovered more held-out in-trial targets than the genetics-only model at every ranking depth, from 5 to 9 genes in the top 50, 11 to 14 in the top 100, 22 to 27 in the top 200, and 57 to 63 in the top 500 (**Figure 4e, Supplementary Table 7**). Within the perturbational layer, residualized trans-regulatory burden contributed the most in the strong genetic prior group, while count of significantly regulated cytokines added the most signal in the weak genetic prior group (**Figure 4f**).

**Figure 4.**
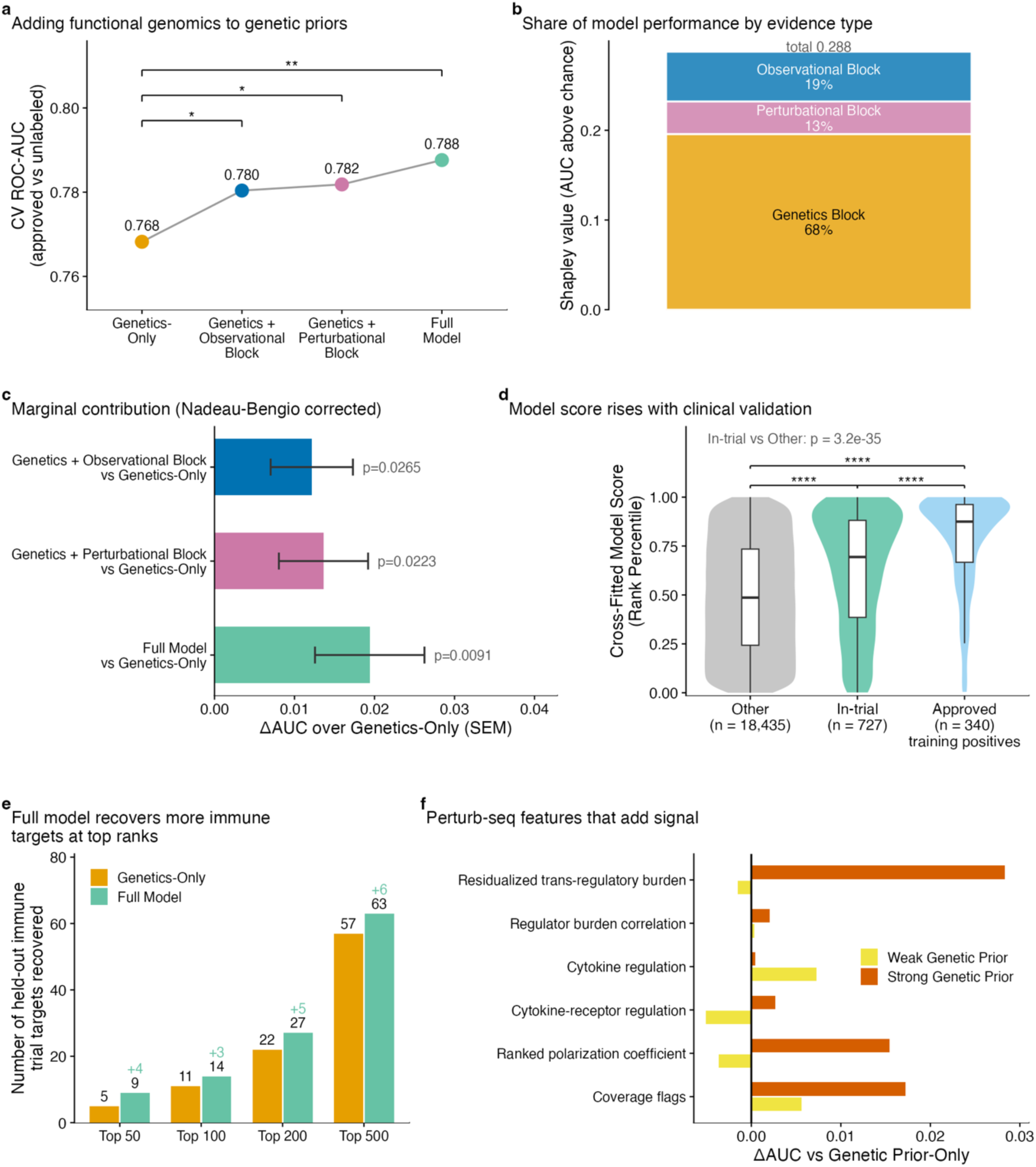
Integrating functional genomics with genetic priors improves recovery of immune targets. **(a)** CV ROC-AUC computed for four nested feature sets, distinguishing held-out approved targets from unlabeled genes. Significance against the genetics-only model was determined by Nadeau–Bengio corrected t-tests. **(b)** Grouped-Shapley decomposition of AUC-above-chance, showing shares attributable to the genetics, observational, and perturbational blocks. **(c)** Marginal AUC increase over genetics-only model (± Nadeau– Bengio corrected SE) for each augmented feature set. *p*-values computed from Nadeau–Bengio corrected t-tests. **(d)** Distribution of cross-fitted model rank percentile across approved, in-trial, and all other genes. Boxplot denotes median and IQR. Brackets show pairwise Mann-Whitney tests, and displayed *p*-value is between in-trial targets and non-targets. **(e)** Number of held-out in-trial targets recovered within top 50, 100, 200, and 500 nominations for the genetics-only and full model. **(f)** Incremental signal contributed by blocks of perturb-seq features, stratified by median split of genetic prior score. * p < 0.05, ** p < 0.01, *** p < 10^−3^, **** p < 10^−4^; ns, not significant.

A separate label-permutation null over 1,000 permutations confirmed baseline calibration across all model variants: the genetics-only model, the genetics and observational functional genomics model, the genetics and perturbational functional genomics model, and the full model. These models were at least 14 standard deviations above their respective null distributions (**Supplementary Figure 1a, Supplementary Table 8**). A scrambled-feature control further confirmed that this performance improvement is driven by true feature signals. Row-shuffling the 19 functional genomics features eliminated the performance increment. 99 scrambled draws resulted in mean AUC gain below zero at −0.0060 ± 0.0035 (SD) against the observed increment of +0.0194, which no draw reached (empirical *p* = 0.010) (**Supplementary Figure 1b**). While the increment itself is small in absolute AUC, this is expected for orthogonal evidence added on top of a strong prior^42^. Functional genomics features added small, orthogonal signals that yielded primary impact at the top of the ranking, where candidate prioritization is most useful. Adding these features increased recovery of held-out in-trial targets by +80% at top 50, +27% at top 100, +23% at top 200, and +11% at top 500.

While gene rankings were generally well correlated between the genetics-only model and the full model across all 19,502 genes (Spearman’s ρ = 0.83), 49 genes inverted across the 25th/75th percentile boundaries. Functional genomics features promoted 36 genes from below the 25th percentile to above the 75th percentile and demoted 13 genes in the reverse direction (**Supplementary Figure 2a, Supplementary Table 9**). The promoted genes interacted with each other significantly more than matched controls and were significantly enriched for a single antiviral gene ontology (GO) term, defense response to virus (**Supplementary Figure 2b, Supplementary Table 9**). Notably, 22.2% of the promoted genes lacked GO biological-process terms compared to 9.5% of coverage-matched non-discordant genes, highlighting the strength of functional genomics in detecting targets that annotation-based prioritization cannot. All nine features that distinguished promoted or demoted genes from matched controls were functional genomics features (**Supplementary Figure 2c**).

### Top model nominations are enriched for targets in immune clinical development

Among the top 50 genes nominated by the full model, after excluding already approved immune-indication targets, nine genes had immune-indicated drugs in trial, with five in Phase II and four in Phase III (**Figure 5a, Supplementary Table 10**). Note that these in-trial targets were held out from the training set and thus did not influence our model parameters. Against bootstrapped controls, the top 50 genes nominated by the full model were 4.8-fold enriched for any immune-trial targets and 5.9-fold enriched for advanced immune-trial targets (**Figure 5b**). The genetics-only model was also enriched over the same background, but less so at 2.7-fold for any-trial and 4.5-fold for advanced targets (**Figure 5b**). Adding functional genomics features improved enrichment for immune trial targets from 4.2-fold to 5.4-fold in the top 25, 2.7-fold to 4.8-fold in the top 50, and 2.9-fold to 3.8-fold in the top 100 nominations (**Figure 5c**).

**Figure 5.**
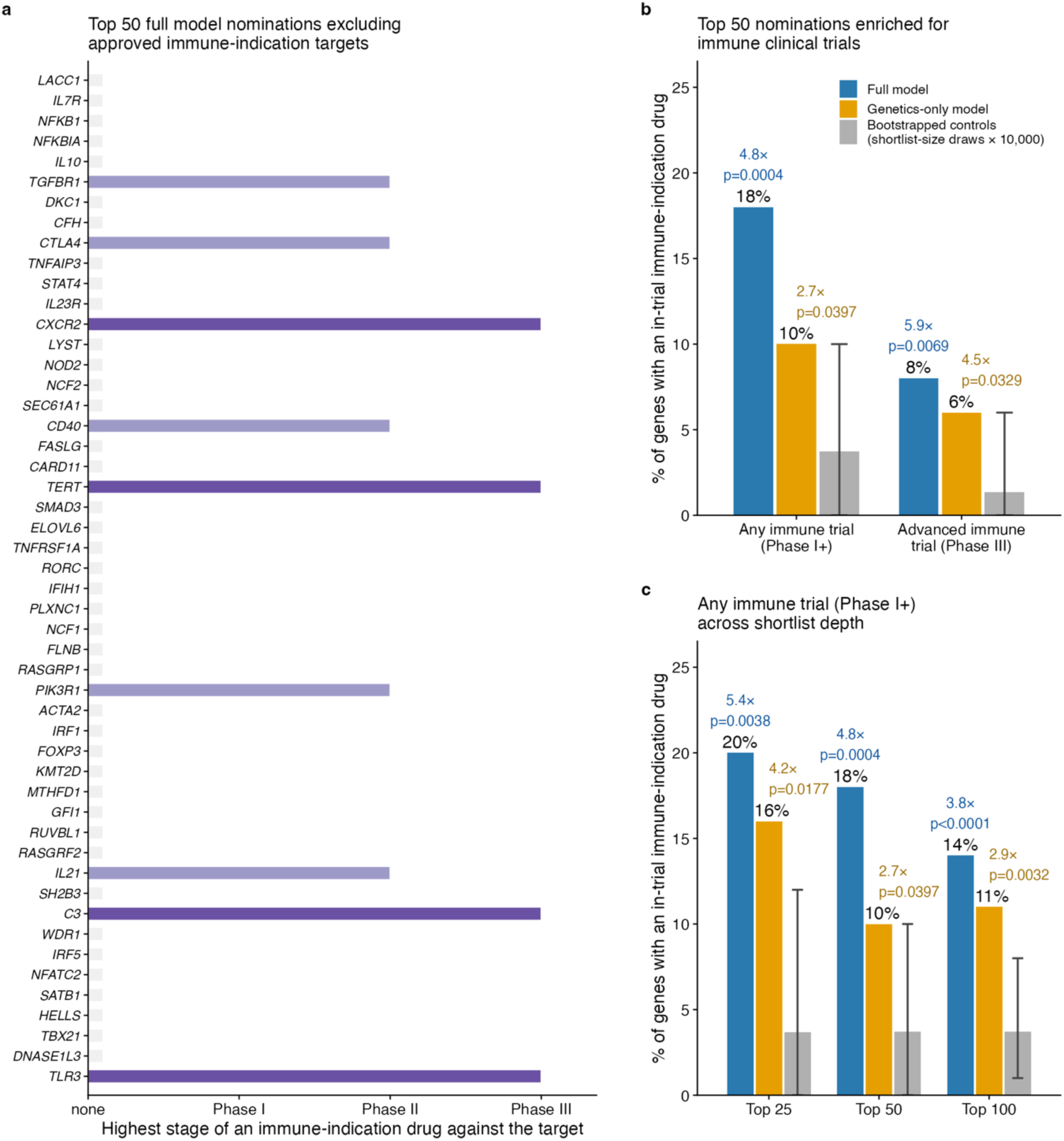
Top nominations are enriched for genes in immune-indication clinical development. **(a)** Top 50 full model nominations after excluding approved immune-indication targets. Genes are ordered by rank and colored by their most advanced immune-indication trial stage. **(b)** Enrichment of the top 50 for genes with any immune-indication trial (Phase ≥ I) and advanced immune trials (Phase III), relative to 10,000 bootstrap draws of 50 control genes sampled from non-approved genes ranked outside the top 200. Colored bars indicate the observed percentage for each model. The gray bars indicate the bootstrap control mean with error bars spanning the 2.5th to 97.5th percentiles of the bootstrap distribution (control mean 3.7% for Phase ≥ I and 1.3% for Phase III). Fold enrichment and adjusted empirical p-values are reported above each model bar. **(c)** Enrichment of genes with any immune-indication trial (Phase ≥ I) across shortlist depths of 25, 50, and 100 for both models. Control draws match the shortlist size, and thus bootstrapped intervals narrow as depth increases while the mean is fixed at 3.7%.

### IGNITE outperforms an external genetic benchmark under leakage control

Genetic Priority Score (GPS) serves as an apt external genetic benchmark because, like IGNITE, it does not rely on gene-level functional annotations such as gene, mammalian phenotype, and human phenotype ontologies or protein-protein interaction (PPI) network propagation as inputs. These annotations are known drivers of performance in drug target prioritization^10,11^ but are also highly vulnerable to coverage bias and confounded signals^13,16,43^. To control for model bias, IGNITE and GPS were evaluated on their recovery of 535 in-trial genes unseen by either model. Restricted to these targets, IGNITE ranked immune-trial targets more accurately, and the DeLong test confirmed that IGNITE AUC was significantly higher than that of the compared model (**Supplementary Figure 3, Supplementary Table 11**). Without leakage control, the two scores were statistically indistinguishable (**Supplementary Figure 3, Supplementary Table 11**).

### Temporal validation confirms IGNITE predicts future clinical advancement

One limitation of cross-sectional labeling is that undiscovered positives in the unlabeled pool are indistinguishable from true negatives^44–46^. This can inflate performance estimates and preclude assessment of whether the model can prospectively anticipate future target discoveries^47,48^.

To address this, we froze labels at *T0* = 2014 and retrained our PU model on the 259 genes approved for an immune indication on or before 2014 (**Figure 6a, see Methods**). This procedure defined 134 emergent genes that were unlabeled at *T0* but have since been approved or in-trial for an immune indication by present-day evidence, and whose first trial of any indication was registered after 2014. Our PU learning model was tested by ranking these 134 emergent genes among the remaining 18,435 unlabeled genes.

**Figure 6.**
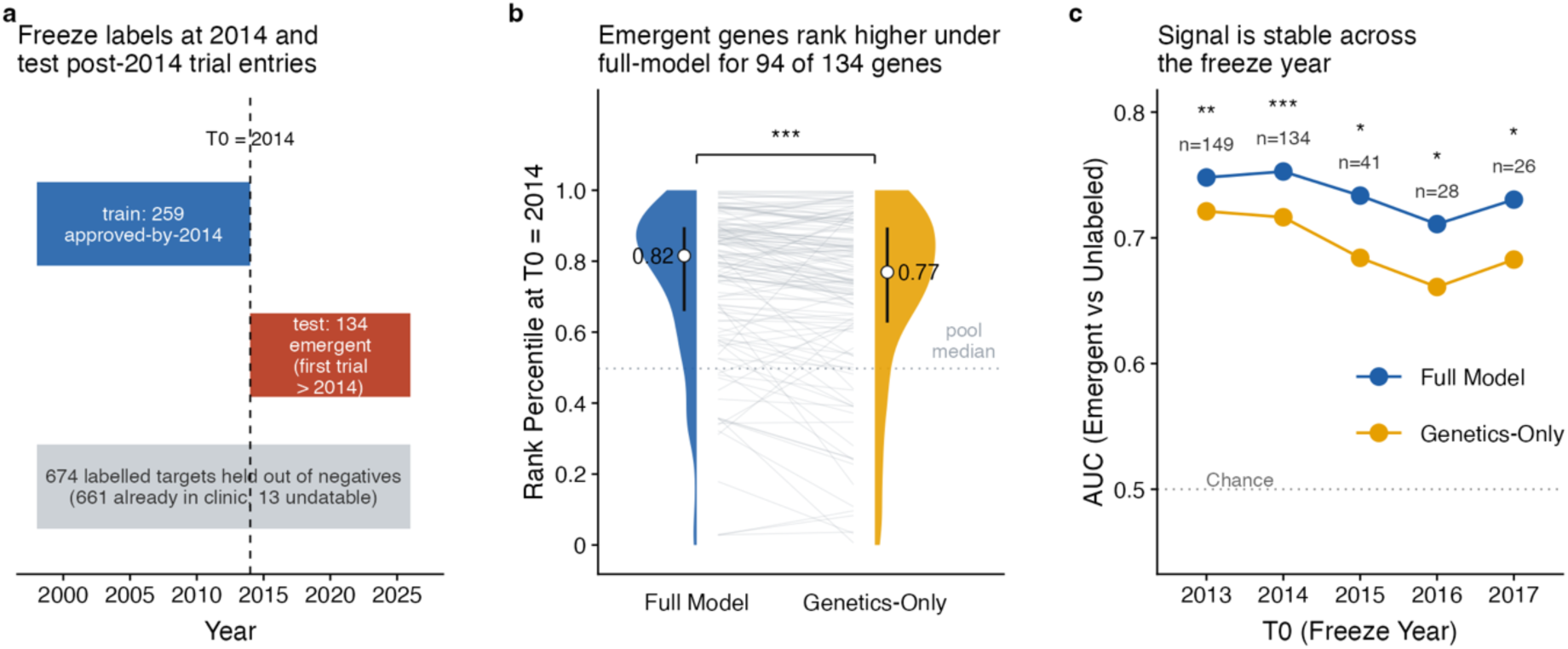
Temporal validation shows IGNITE predicts genes that entered immune trials after label freeze at *T0*. **(a)** Design of temporal holdout (see Methods). **(b)** Rank percentiles of 134 emergent genes at *T0* = 2014 under the full model and genetics-only model, shown as half violin plots with lines connecting each gene across models. Points and bars denote each model’s median and IQR. The dotted line denotes the median rank of the unlabeled genome. Significance was assessed by comparing the AUCs for emergent versus unlabeled genes between the two models using the paired DeLong test. **(c)** Freeze-year sensitivity analysis comparing the genetics-only model and full model with *T0* varied from 2013 to 2017. Significance was assessed by the paired DeLong test at each freeze year. * p < 0.05, ** p < 0.01, *** p < 10^−3^, **** p < 10^−4^; ns, not significant.

The full model, which includes functional genomics features, outperformed the genetics-only model in ranking emergent genes above the unlabeled genome (**Figure 6b, Supplementary Table 12**). This effect was also stable across freeze years, with the full model significantly outperforming the genetics-only model at all freeze years (**Figure 6c, see Methods**).

### IGNITE’s target recovery is immune-specific

IGNITE preferentially recovered immune-exclusive targets, suggesting that it encodes immune-specific biology rather than generic druggability signatures (**Figure 7**). On immune-exclusive targets, IGNITE significantly outperformed the indication-matched genetic comparator and the off-indication genetic control (**Figure 7a, Supplementary Table 13**). On cardiac-exclusive targets, IGNITE performed significantly worse than the indication-matched cardiac genetic control and comparably to the immune genetic comparator (**Figure 7b**).

**Figure 7.**
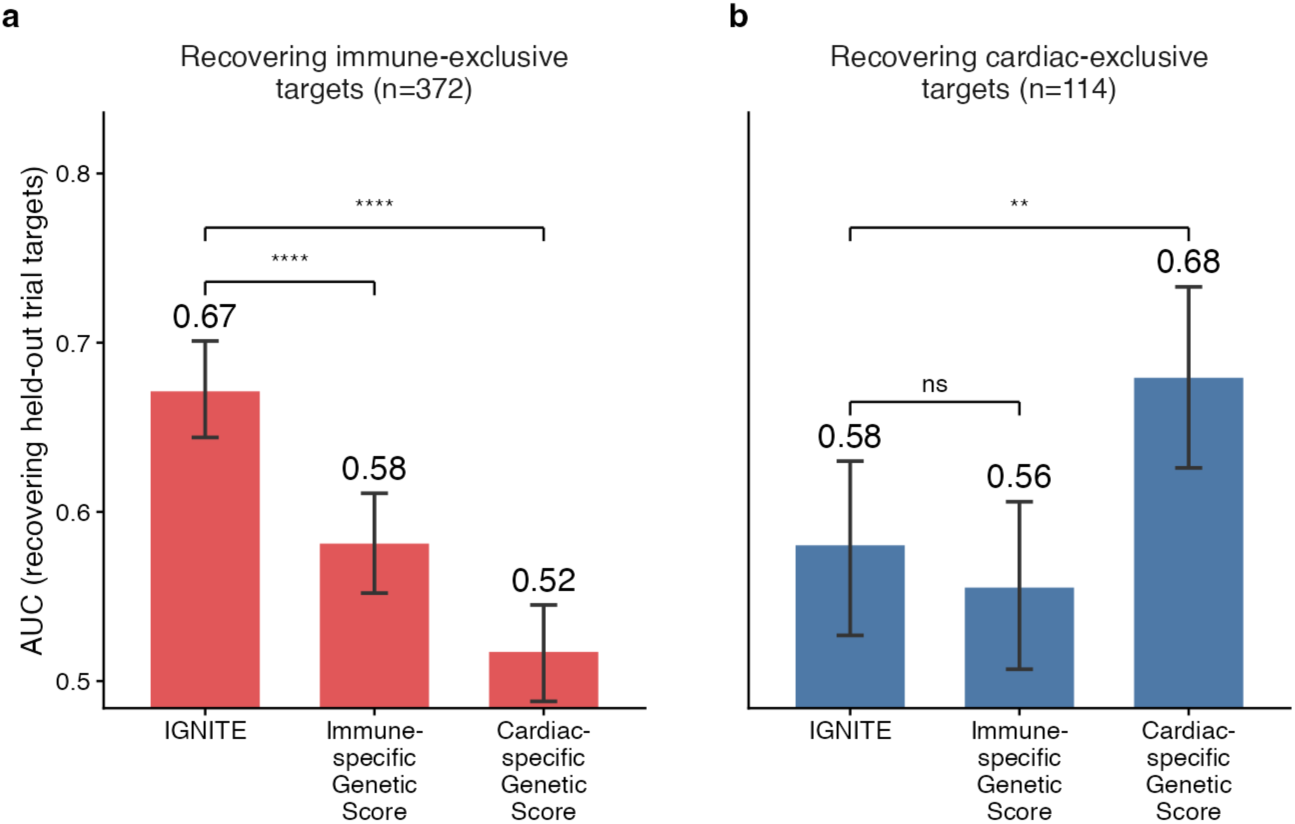
Target recovery by IGNITE is immune-specific. **(a)** Recovery of immune-exclusive targets by IGNITE, the Open Targets immune-genetic, and the Open Targets cardiac-genetic scores. Quantified as ROC-AUC (±95% CI) for ranking held-out trial targets. Error bars are derived from 2,000 bootstrap draws. Significance assessed by paired DeLong test. **(b)** Recovery of cardiac-exclusive targets by the same three scores. Significance assessed by paired DeLong test. * p < 0.05, ** p < 0.01, *** p < 10^−3^, **** p < 10^−4^; ns, not significant.

### *STAT4* is a validated immunoregulator that remains undrugged

*STAT4* is a compelling example of a top-ranked gene nominated by IGNITE that currently has no selective drugs approved or in trial for any indication^39,49^ (**Figure 8**). STAT4 activity can be targeted pharmacologically, as its activity can be modulated indirectly through upstream inhibition of JAK. Direct human evidence for this exists where ruxolitinib reduced dermatologic and inflammatory phenotypes in patients with *STAT4* gain-of-function (GOF) syndrome^50^. The first *STAT4*-selective small molecules were also reported only recently and remain preclinical^51^. As part of the JAK-STAT pathway, *STAT4* is a long-recognized immunoregulator associated with multiple autoimmune and inflammatory diseases^50,52,53^.

**Figure 8.**
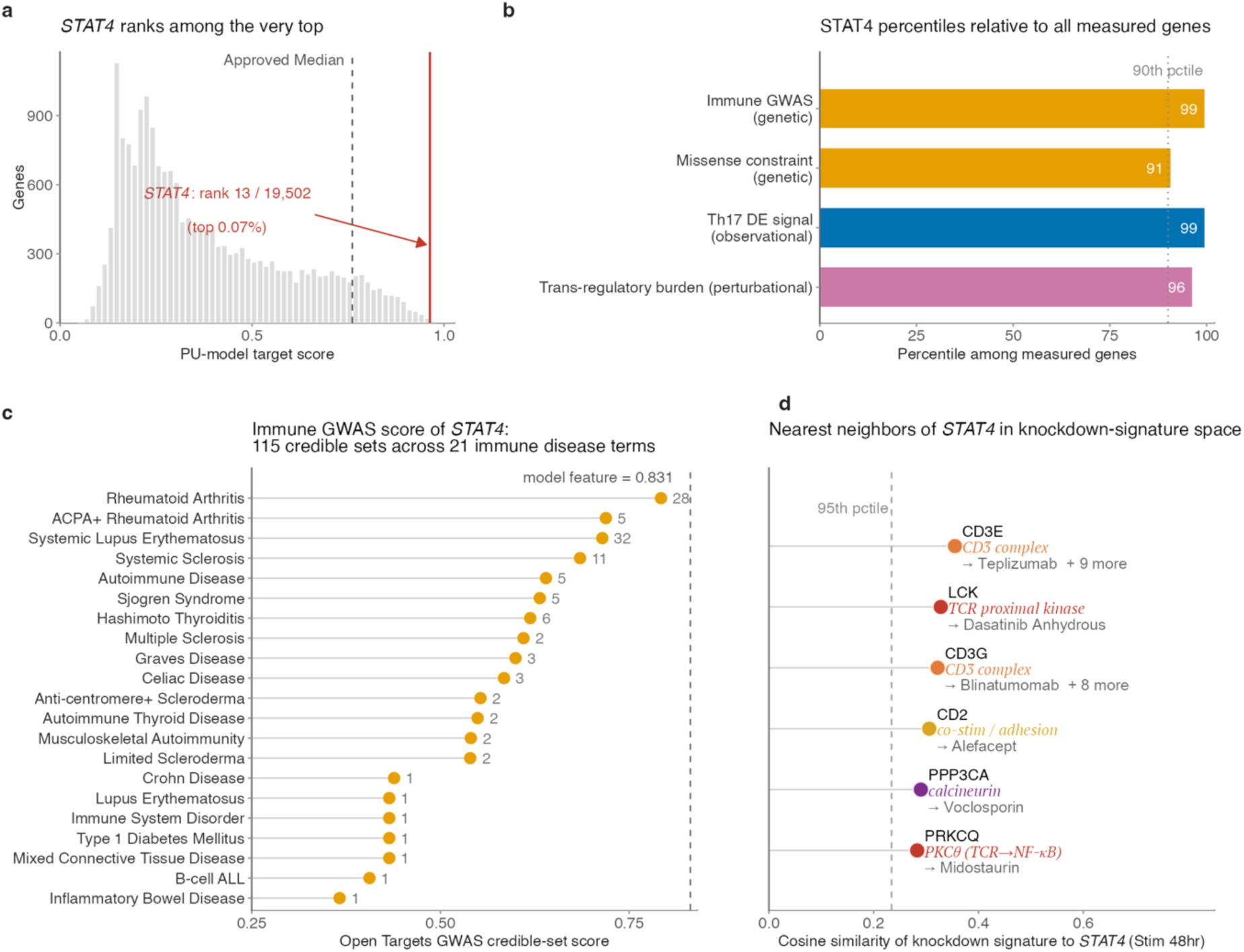
IGNITE nominates *STAT4*, a genetically and functionally supported but undrugged target. **(a)** Distribution of PU-model target scores across the genome, with *STAT4* and the approved-target median indicated. **(b)** *STAT4* percentiles relative to all measured genes on four evidence axes. **(c)** Immune disease GWAS score for *STAT4* is broken down into 21 contributing immune disease terms and ordered by their credible-set association scores. Numbers next to each point indicate the number of credible sets per disease term. The dashed line denotes the composite immune indirect GWAS score used in IGNITE. **(d)** Cosine similarity of *STAT4* 48hr-stimulation knockdown signatures to those of its nearest approved immune target neighbors, annotated with representative approved drugs. The dashed line denotes the 95th percentile of STAT4’s cosine similarity across all 7,044 profiled genes that are not approved targets.

IGNITE ranked *STAT4* 13th out of 19,502 genes with a probability score of 0.963, placing it far above the approved target median of 0.761 (**Figure 8a, Supplementary Tables 1 and 14**). The candidate was also supported by four types of evidence: an immune disease GWAS score at the 99th percentile, Missense Z-score at the 91st, Th17 differential expression at the 99th, and residualized trans-regulatory burden at the 96th (**Figure 8b**). Decomposition of the immune disease GWAS score revealed that *STAT4* harbors non-coding common-variant associations across a broad range of immune-mediated disorders including rheumatoid arthritis, systemic lupus erythematosus, and systemic sclerosis (**Figure 8c, Supplementary Table 14**). Beyond these common variants, *STAT4* also carries rare coding variants at both extremes, a germline GOF allele that causes disabling pansclerotic morphea and LOF alleles underlying its IEI inclusion^50^. Among approved immune targets, the *STAT4* knockdown signature was most similar, by cosine similarity, to those of genes associated with T-cell activation, including *CD3E*, *LCK*, and *CD3G*, consistent with the role of the JAK-STAT pathway in T-cell activation, survival, and differentiation (**Figure 8d**).

While *STATs* have long been considered undruggable, that view is shifting^54–56^. This trend is exemplified by growing interest in *STAT3*, which ranks in the top 1.3% in IGNITE^57–60^. We therefore highlight *STAT4* as a compelling example of a drug target candidate nominated by IGNITE. STAT4 is likely to become more tractable as small-molecule binders and biologics improve.

### *ELOVL6* and *RUVBL1* are novel candidates nominated by IGNITE

While *STAT4* is already a well-supported drug target by multiple lines of evidence apart from IGNITE, our model’s greater value lies in surfacing candidates missed by genetics-only and annotation-based prioritization methods. Two candidates from the top 50 nominations, *ELOVL6* and *RUVBL1*, are interesting examples of novel candidates whose canonical functions have no apparent immune relevance. Moreover, neither gene is associated with Mendelian immunodeficiency or has strong GWAS or gene burden signals for immune disease. Adding the functional genomics layer boosted the rankings of both genes from outside the top 1,000 into the top 50. *ELOVL6* rose from rank 1,549 in the genetics-only model to 27 in the full model, and *RUVBL1* rose from 1,240 to 47.

ELOVL6 is the primary enzyme that elongates C16 palmitate to C18 stearate within the mammalian endoplasmic reticulum, a canonical role in lipid metabolism with no apparent immune relevance^61^. However, it is positioned immediately downstream of the ACC1-FASN de novo lipogenesis cascade, a pathway that is required for Th17 differentiation and whose inhibition diverts CD4^+^ T cells towards FOXP3^+^ Tregs and attenuates experimental autoimmune encephalomyelitis^62–65^. The experimental immune evidence for ELOVL6 involves multiple systems including the airway, CNS, liver, and hematopoietic system^66–69^. Most importantly, ELOVL6 is already highly tractable with multiple potent, orally active, selective inhibitors^70–74^.

RUVBL1 is a chromatin-associated AAA+ ATPase whose canonical roles in chromatin remodeling, transcription, and DNA repair have no apparent immune relevance^75,76^. However, RUVBL1 drives transcription of Type I interferon-stimulated genes (ISGs) by specifically binding STAT2 and recruiting RNA pol II to ISG promoters^77,78^. Clinically, anti-RuvBL1/2 autoantibodies are linked to a specific overlap syndrome of systemic sclerosis and myositis^79,80^. Most importantly, in contrast to most chromatin scaffold proteins, RUVBL1 is pharmacologically tractable with a preclinical allosteric inhibitor that is effective in vivo^81,82^.

## Discussion

While human genetic evidence is now a standard input to target selection^1,6^, current genetic features are largely associational and thus prioritize genes whose variation tracks disease risk without accounting for how they function in the disease-relevant human cell state. Here, we evaluated the impact of observational and perturbational functional genomics on target prioritization using the largest publicly available genome-scale perturb-seq dataset in primary human cells to date. Because this resource is derived from primary human CD4^+^ T cells, we focus on immune indications.

We demonstrate that both observational and perturbational functional genomics in the relevant primary cell type carry target-relevant information that genetics alone does not. Adding functional genomics to the genetics-only model improved cross-validated AUC significantly, although by a modest margin, which is expected when orthogonal evidence is added to an already strong genetic prior. Importantly, this integrated model outperformed an indication-matched genetic control on immune-exclusive targets but underperformed a cardiac comparator on cardiac-exclusive targets, suggesting that the model learned cell-type specific features relevant for immune diseases.

The five genetic priors are informative because they span four non-overlapping evidence types: rare high-penetrance coding evidence in IEI, common variant associations in immune disease GWAS scores, rare variant associations in immune disease gene burden scores, and disease-agnostic genic intolerance metrics in Missense Z-score and LOEUF. The functional genomics layer provides an orthogonal axis to these priors. Notably, residualized trans-regulatory burden nominates targets with strong genetic priors while regulated cytokine count nominates targets with weak genetic priors. The practical value of the functional genomics layer lies at the top of the rankings, where recovery of held-out in-trial targets improved by 80% at the top 50 nominations and by 27% at the top 100. A scrambled-feature control indicates that these gains reflect genuine signal rather than added model capacity. In temporal validation, IGNITE also outperformed the genetics-only model in predicting genes that would subsequently enter immune trials.

Most strikingly, functional genomics also reaches genes that annotation-based evidence cannot. Among the genes promoted by the functional genomics layer, 22.2% lacked GO biological-process terms compared to 9.5% of coverage-matched genes, and every feature that distinguished promoted genes from matched controls was a functional genomics feature. The value of functional genomics in target prioritization is further demonstrated by our two novel candidates *ELOVL6* and *RUVBL1*, which moved from outside the top 1,000 into the top 50.

We intentionally did not train IGNITE on druggability features. This design choice prioritizes biological evidence to avoid anchoring to tractability assessments that both confound genetic signal^83^ and shift as small-molecule binders and biologics improve^84,85^. *STAT4*, one of our candidates, illustrates this rationale. STATs were historically regarded as undruggable, an assessment that is now changing, as shown by the trajectory of STAT3^54–56^ and recently reported preclinical *STAT4*-selective molecules^51^.

Several limitations should be considered. First, positive-unlabeled learning treats unlabeled genes as provisional negatives when some are undiscovered targets, which results in conservatively lower performance estimates. Second, all functional genomics features are derived from a single primary CD4^+^ T-cell dataset and do not include other immune lineages such as CD8^+^ T cells, B cells, and myeloid cells. However, this reflects the current availability of genome-scale perturb-seq data rather than a design choice, and the framework can extend without modification as these resources become available. Third, the label independence evaluated in this study is temporal rather than cross-database. Because most publicly available annotation sources rely on ChEMBL, substituting an annotation source would not change the data provenance. Moreover, even an independently curated set would largely converge because approved-target status is a public record, making temporal validation the most informative test of independence for this study. Fourth, while temporal holdout is an effective independence test, it is not truly prospective due to data limitations. Although the labels are frozen at *T0*, the features themselves are not frozen in time and derive from present-day resources. Since the model has access to feature information that did not exist at *T0*, the holdout test only approximates and does not fully reproduce a true prospective analysis. Fifth, genetic priors are largely derived from resources that lack ancestral diversity, which limits resolution of gene-level estimates such as intolerance^5,86^.

In summary, we have developed IGNITE to prioritize gene targets for immune-mediated disease by integrating observational and perturbational functional genomics with human-genetic priors. IGNITE has utility across the immune drug development pipeline from surfacing novel candidates to corroborating targets already in clinical trials, with its greatest value concentrated at the top of the ranking. As the number of approved immune drugs grows and genome-scale perturbation data extend to additional immune lineages and stimulation contexts, we expect IGNITE’s performance to improve further. More broadly, our work adds to the growing evidence that experimental readouts of gene function in the disease-relevant human cell type carry target-relevant information beyond human genetics, and that combining functional genomics with genetic priors is a productive route to more accurate, indication-specific target prioritization.

## Methods

### Gene universe and drug-status labels

We defined the model universe as 19,502 protein-coding genes and obtained drug-target annotations from the Open Targets Platform 26.06. Each gene was labeled with the most advanced clinical stage reached by any drug that targets it in an immune indication, defining immune indication as the immune system disorder subtree of the Mondo Disease Ontology (MONDO:0005046 and its descendants). Clinical stage was determined at the indication level rather than per drug. Because Open Targets annotates mechanism of action targets at the drug level, each target of a multi-target drug received equal credit, and multi-target drugs were expanded into their constituent targets. Genes whose furthest-stage annotation was unknown were treated as non-targets. Of the 734 in-trial genes, 7 were mitochondrially encoded and absent from the nuclear feature matrix, leaving 727 in-trial targets within our feature matrix. This resulted in our final 19,502-gene feature matrix consisting of 340 approved targets, 727 in-trial targets, and 18,435 non-targets. The reported *n* is the number of genes with data for a given assay and thus varies with each assay’s gene coverage.

### Feature construction

The feature matrix consisted of 24 features in three evidence blocks: human genetics and genic intolerance, observational functional genomics, and perturbational functional genomics. Block membership was defined once and asserted at each attribution-decomposition step to cover all non-metadata columns exactly, ensuring every modeled feature enters decomposition (**Supplementary Table 2**). The first block captured human genetics and genic intolerance and comprised IEI inclusion, immune disease GWAS score, immune disease gene burden score, Missense Z-score, and LOEUF. IEI inclusion was encoded as a binary variable from the International Union of Immunological Societies (IUIS) classification^38^. We derived the immune system disorder indirect GWAS and burden scores from Open Targets 26.06 using MONDO:0005046 and its descendants^39^. Genes without GWAS or burden scores were assigned zero. Both Missense Z-score and LOEUF were derived from gnomAD^40^. Both observational and perturbational functional genomics blocks were derived from a common CD4^+^ T-cell resource^32^. Observational features include gene-level differential expression z-scores for polarized Th1, Th2, Th17, and Treg cells relative to resting Th0 cells. Perturb-seq features include gene-level ranked polarization coefficient, regulator burden correlation, number of regulated cytokines, and number of regulated cytokine receptors. Each of these was measured at rest and in the 8hr- and 48hr-stimulated conditions. Here, we transformed regulator burden correlation into a binary variable encoding significance of LOF burden on lymphocyte count. In addition, we summarized the range of ranked polarization coefficients across conditions, and we estimated residualized trans-regulatory burden based on established methods^32^. A binary coverage indicator was added to flag genes for which cytokine and cytokine receptor data were available. Missing values were imputed with training-row medians, which were computed separately within each bagging iteration. This prevented test-fold information from entering the imputation. Columns that were entirely missing within a given fit were set to zero.

### Positive-unlabeled learning model

Genes were prioritized using positive-unlabeled bagging. Over T iterations, a pseudo-negative set of size equal to the number of training positives was sampled with replacement from the unlabeled genes, and a gradient-boosted tree classifier (XGBoost v3.3.0; 120 trees, max depth 2, learning rate 0.1, subsample and column sample 0.8) was fit against the approved positives. Each gene’s score was its mean probability across the iterations in which it was not sampled as a pseudo-negative:

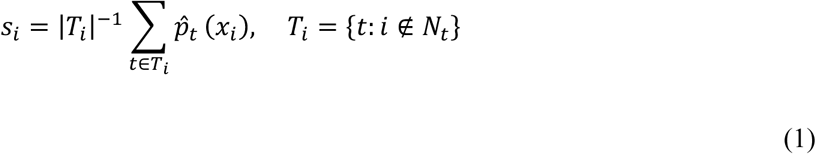

where *s_i_* is the final ensembled score for item *i*. |*T_i_*| is the number of times item *i* was out-of-bag. *t* is the index for outer bagging iteration of the complete XGBoost model. *p̂_t_*(*x_i_*) is the predicted probability from the model at iteration *t*. *N_t_* is the subset of pseudo-negative samples for training at iteration *t*.

Pseudo-negatives came only from the unlabeled pool, so the 727 in-trial genes never entered training and were scored on all T iterations, a clean holdout. An unlabeled gene, by contrast, served as a pseudo-negative whenever it was drawn, and those rounds were excluded from its own score. In the single-fit configurations, the approved positives were in the training set of every iteration and thus scored in-bag. Those scores were reported for complete ranking but were not used in any performance estimate. Under cross-validation, each fold’s held-out positives were scored out-of-bag. For the drug-status group comparison (**Figure 4d, Supplementary Table 7**), approved positives were split into five folds, and the model was trained at *T* = 200 on the remaining positives in each fold. Each non-positive gene was scored by cross-fitting, averaging across folds that did not train on it, while positives were held out once and scored on all iterations of that fold’s fit. This placed all three groups on a common out-of-sample basis. The published ranking retains the single-fit score.

### Model configurations

Four configurations of this model were used for performance estimation and attribution. The fifth, described above, was used to produce cross-fitted scores for drug-status comparison (**Figure 4d**). First, the ranked atlas was a single fit on all 340 positives at *T* = 200 and the canonical seed, without cross-validation. Second, cross-validated performance used five folds across 10 seeded repeats at *T* = 40. Third, the evidence-coalition ladder used 5 folds across 5 seeds, giving 25 AUC samples per coalition at *T* = 30, because it fits all 8 coalitions of the three blocks. In the ladder and in the per-block stratum analysis, the base learner is seeded by bagging iteration alone whereas the atlas and repeated-CV fits offset the iteration by the run seed.

In both cross-validated configurations each seed sets the cross-validation partition, and within a seed the pseudo-negative draws are common to the 5 folds. Folds were taken over the positives alone, so the fold generator received a single class label, and no class stratification occurred, leaving a shuffled equal-sized partition of the positives.

The per-block stratum analysis is the fourth configuration. Here, genes were split into weak- and strong-prior strata at the median genetics-only prior score of unlabeled genes at *T* = 100 and the canonical seed. Within each stratum, the signal contributed by a perturb-seq feature group was defined as AUC(prior + group) − AUC(prior) where AUC measures discrimination of held-out in-trial genes from unlabeled genes at *T* = 60. For the prior model and each prior + group model, out-of-bag scores were computed for all 19,502 genes and averaged across five seeds. A single AUC was then computed per stratum from the averaged scores. Perturbation-derived features were partitioned into six groups and evaluated: trans-regulatory residual, regulatory burden, cytokine regulation, cytokine-receptor regulation, ranked polarization coefficients, and coverage flag. The observational functional genomics features were excluded by design.

### Attribution

Average attribution quantifies contribution of each of the three predictor blocks to discrimination through a grouped Shapley decomposition of AUC-above-chance. The value of coalition *S* is:

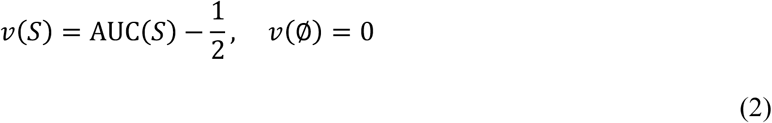

where *v*(*S*) is the value assigned to a specific coalition of predictors *S*. AUC(*S*) is the AUC-ROC achieved by a model trained exclusively on features within coalition *S*. ∅ represents empty coalition and thus a model with no predictors.

The exact Shapley value of each block is:

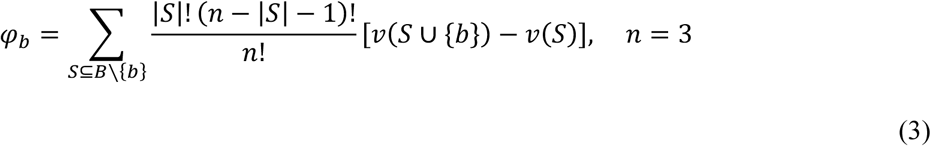

where *φ_b_* represents the average marginal contribution of predictor block *b*. *B* is the complete set of all predictor blocks. *S* is the subset of predictor blocks without block *b*. |*S*| is the number of predictor blocks currently in subset *S*. *n* is the total number of predictor blocks, which is 3 in our case. *v*(*S* ∪ {*b*}) − *v*(*S*) measures the marginal increase in predictive performance when block *b* is added to coalition *S*.

All eight coalitions are enumerated directly, avoiding the need for sampling approximations or external libraries. Furthermore, Shapley values are computed at the block level and do not scale with the number of features in each.

Marginal attribution is the paired across-fold ΔAUC over the genetics-only coalition. Nadeau–Bengio corrected t-test was used to test significance.

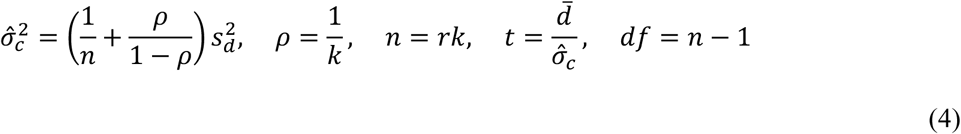

where 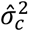 is the corrected estimator for performance differences adjusting for overlapping training data. *n* is the number of total folds evaluated. *ρ* is the proportion of the total dataset held out as the test set in each fold. 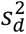 is the standard sample variance of performance differences across all evaluated folds. *k* is the number of folds in the *k*-fold cross-validation, and *r* is the number of times the *k*-fold cross-validation is repeated. *t* is the test statistic. *d̅* is the average difference in AUC across all folds and *σ̂_c_* is the corrected standard error. *df* is the degrees of freedom for the t-distribution.

### Null models

Two nulls were computed. First, a label-permutation null over 1,000 permutations reassigned 340 positive labels at random, without replacement, from the approved and unlabeled gene set excluding the 727 in-trial genes. For each permutation, the four-rung cross-validated AUC ladder was re-run across five seeds and five folds, calibrating the level of each of the four rungs. Second, a scrambled-feature control calibrated the increment by holding true labels and true genetic features fixed while row-permuting the functional genomics features. This preserved each feature’s marginal distribution and model capacity while removing gene-to-row correspondence and covariance among functional genomics features. 99 total draws were performed.

### Trial-target recovery and enrichment control

Model performance was assessed with two complementary metrics. Global discrimination was quantified as ROC-AUC, indicating whether held-out in-trial targets rank above the unlabeled genome. Because the utility of a prioritization score is largely defined by its discrimination ability at the shortlist level, fold enrichment and top-*k* recovery are reported as performance estimates. Fold enrichment was computed at shortlist depths of 25, 50, and 100 with a fixed control pool drawn from beyond rank 200. Held-out recovery counted the number of held-out in-trial genes among the top-*k* of the model’s ranking of the 19,162 non-approved genes, at k = 50, 100, 200, and 500. The trial stage was derived from Open Targets Platform 26.06, and thresholds were applied where Phase ≥ I for any immune trial and Phase ≥ III for any advanced immune trial. For the bootstrapped enrichment analysis, each model drew 10,000 control sets equal in size to the shortlist, sampling with replacement from the non-approved genes ranked outside that model’s top 200 nominations. Fold enrichment was defined as the observed shortlist rate divided by the mean control rate. The empirical *p*-value was computed as the fraction of control draws reaching or exceeding the observed shortlist rate and was then adjusted by the Benjamini–Hochberg false discovery rate. A paired DeLong test was used to compare the full model against the genetics-only model on predicting held-out in-trial genes. This was a single test and is thus reported uncorrected.

### Comparison against published Genetic Priority Score (GPS)

Because GPS reports multiple indication-specific scores per gene, we used the maximum score per gene to compare against IGNITE. GPS covered 14,677 of the 19,502 genes, and thus genes without a GPS score were assigned zero. We first evaluated the recovery of all 727 trial targets without removing genes that GPS had used as positive drug labels during training. GPS used 453 positive drug labels during training. Of these, 284 fell inside our non-approved pool, 161 were in our immune approved pool already, and eight were absent from our gene universe. To prevent data leakage, we then excluded 284 of those genes from the evaluation, which removed 192 positive genes from the held-out in-trial set and 92 unlabeled genes. This left 535 held-out in-trial targets within an 18,878 gene universe that neither GPS nor IGNITE had trained on as positives, enabling a leakage-controlled comparison. Paired DeLong tests were used in evaluating IGNITE against GPS on predicting leakage-controlled test sets. Comparison between IGNITE and GPS was run on standard and leakage-controlled evaluation sets and corrected together by the Holm– Bonferroni method.

### Temporal validation

Labels were frozen at a cutoff year *T0*, and the PU model was retrained using only genes approved for an immune indication on or before *T0*. It was then evaluated on genes whose first trial for immune indications started after *T0*. For each gene, the trial date was defined as the earliest start date among all drugs annotated to that gene. At the headline freeze year *T0* = 2014, 259 genes had been approved for an immune indication and formed the training positives. Of the remaining 808 approved or in-trial targets, 661 genes had already entered clinical trials or gained approval for any indication by 2014. A further 13 genes lacked both a datable trial and an approval. Because these genes are established drug targets by the freeze year or cannot be positioned on the timeline, they were excluded from the pseudo-negative pool and emergent set. This left 134 emergent genes that were unlabeled at *T0* but are approved or in-trial for an immune indication by present-day evidence, with a first trial of any indication registered after 2014 (**Supplementary Table 12**). We tested whether our PU learning model could rank these 134 unlabeled emergent genes among the other 18,435 unlabeled genes.

Here, both the pseudo-negative pool and emergent test set exclude any labeled target gene with pre-*T0* evidence of trial or approval in any indication. This constraint minimizes the probability that an emergent gene’s post-*T0* trial entry reflects clinical relevance the model already had access to through an unrelated developmental pathway. Sensitivity was assessed over freeze years 2013 to 2017, and 2018 was excluded as underpowered with only 18 emergent genes. Of note, because features reflect contemporary evidence, our analysis is a prospective test of label recovery rather than a fully prospective test that reconstructs the evidence base from *T0*. Paired DeLong tests were used to compare discrimination of emergent genes between full and genetics-only models. Each freeze year was also tested against a permutation null (10,000 draws at headline freeze year *T0* = 2014 and 5,000 draws across the sensitivity sweep). The sensitivity analysis is a sweep of a single pre-specified comparison rather than five independent hypotheses and is thus reported without multiple-testing correction.

### Immune-specificity recovery sets

Immune- and cardiac-exclusive recovery sets were built using Open Targets Platform 26.06 by first filtering for immune-indicated and cardiovascular-indicated drugs with approval or in trial. A gene was considered immune-exclusive if it was in trial for an immune indication and not approved or in trial for a cardiac indication. The converse applied for cardiac-exclusive genes. The shared negative pool was genes unlabeled in both indication areas. Approved immune targets were excluded from both immune- and cardiac-exclusive evaluation sets because IGNITE was trained on them and recovering them would not test specificity. Each recovery set was evaluated by ROC-AUC with intervals from 2,000 bootstrap draws. DeLong test was used to compare models within the same mutually exclusive test set. Pairwise comparisons within each recovery set were corrected for multiple testing using the Holm–Bonferroni method.

### Genetics-versus-full discordance

Discordance was defined on two ranked percentile vectors from the genetics-only and the full model. Promoted genes are below the 25th percentile under the genetics-only model but rise above the 75th percentile under the full model. Demoted genes are above the 75th percentile under the genetics-only model but fall below the 25th percentile under the full model.

Promoted and demoted sets were each compared against control genes sampled from non-discordant genes, which were matched on functional genomics coverage and genetics-only percentile decile to test difference in protein-interaction connectivity and individual functional genomics features. For the feature comparison, 20 matched sets were drawn and pooled into a single control group by direction. Individual features were compared by two-sided Mann-Whitney U test with rank-biserial correlation effect sizes. Benjamini– Hochberg correction was applied within each direction. Protein-interaction connectivity was quantified as the number of STRING interactions among genes in each set and compared against 500 matched null gene sets, with a one-sided empirical *p*-value. STRING interactions were obtained from v12.0 and restricted to edges with a combined score equal to or greater than 400.

GO enrichment used one-sided Fisher’s exact tests over biological-process terms with Benjamini– Hochberg correction applied separately within each direction and each of two backgrounds: all non-discordant genes, and non-discordant genes with both observational and perturbational functional genomics coverage. The correction family was fixed in advance as every GO term annotated to 10-500 genes across the foreground and background. The filter used term size only and was thus independent of a term’s foreground count. Under the primary background, families were 1,410 terms for promoted and 1,409 for demoted, with significance defined at FDR < 0.05. Terms were tabulated if at least three foreground genes carried them. The coverage-restricted background was primary, and the unrestricted background served as sensitivity analysis.

### Knockdown-signature neighborhood

The candidate’s transcriptome-wide knockdown signature was scored by its cosine similarity to the knockdown signatures of each approved target in the resting, 8hr-, and 48hr-stimulated conditions, creating one signature matrix per condition over the 10,282 measured genes (**Supplementary Table 15**). Signatures were perturb-seq Z-scores, and each gene was scored by the maximum cosine similarity to any approved target anchor after zeroing its own knockdown column (112 of 340 approved targets in the stimulated conditions and 105 at rest). In the 48hr-stimulated condition, the comparison distribution consisted of 7,044 genes with an available signature, which included 6,762 unlabeled genes, 223 in-trial genes, and 59 genes outside the feature matrix (**Supplementary Table 14**).

### Statistical analysis

Discrimination was quantified as the ROC-AUC for distinguishing out-of-fold approved genes from unlabeled genes, using five-fold cross-validation across 10 seeds at *T* = 40. All eight coalitions of the three evidence blocks were evaluated by five-fold cross-validation averaged over five seeds, with the empty coalition fixed at chance-level performance. Coalition standard deviations were computed as population standard deviations across the resulting 25 samples.

Model performance was compared across cross-validation folds for each nested feature coalition using the Nadeau–Bengio corrected resampled paired t-test (**Equation 4**). When two scoring methods were evaluated on the same gene panel, they were compared by paired DeLong tests. When gene panels were not shared, the difference was assessed by bootstrapping.

Multiple-testing correction was applied within each pre-specified family of tests rather than across the study as a whole, and the correction applied to each family is stated with the analysis it belongs to. The Holm– Bonferroni method was used for families of five or fewer pairwise comparisons. Benjamini–Hochberg correction was used to screen across many features.

The marginal contributions of each evidence coalition over the genetics-only baseline are reported without multiplicity correction because the three comparisons are nested against a shared baseline model rather than being independent hypotheses, and the Nadeau–Bengio variance inflation already accounts for dependence across cross-validation folds. Permutation nulls and descriptive feature-orthogonality correlations are reported uncorrected.

Empirical *p*-values were computed using two estimators. Most resampling tests counted the fraction of draws that matched or exceeded the observed statistic using the continuity-corrected form. These included the label permutation, scrambled-feature control, temporal-holdout permutation null, and the STRING interaction test. The trial-enrichment analysis reported the fraction of control draws matching or exceeding the observed shortlist rate without the continuity correction.

Enrichment of individual features by drug status was tested with two-sided Fisher exact tests, which reported sample odds ratios with Woolf 95% confidence intervals, and Holm–Bonferroni correction was applied. Kruskal–Wallis tests assessed whether cross-fitted PU score distributions differed across approved, in-trial, and non-target genes, followed by post hoc pairwise Mann–Whitney U tests corrected by the Holm– Bonferroni method.

## Use of large language models

Parts of the analysis code were implemented with the assistance of AI coding agents (Claude Code and Claude Science, Anthropic; models Claude Opus 5.0 and Claude Sonnet 5.0). These tools were used to translate author-written analysis specifications into Python and R, curate databases via API calls, generate plotting code, write documentation, and debug. All prompts were written by the authors, and the choice of methods, parameters, and thresholds was made by the authors, not the AI agents. All AI-generated code was read, tested, and edited by the authors, and all reported outputs were verified by the authors. The analysis code required to reproduce every reported figure and statistic is deposited as described under Code availability.

## Data Availability

Genome-scale perturb-seq in primary human CD4^+^ T cells data are available at the following URL: https://virtualcellmodels.cziscience.com/dataset/genome-scale-tcell-perturb-seq

T-helper polarization differential expression data are available at the following URL: https://github.com/emdann/GWT_perturbseq_analysis_2025

Immune system disorder GWAS and burden scores (MONDO:0005046); Genetic-association scores for immune and cardiac indications are available at the following URL: (Open Targets 26.06) https://platform.opentargets.org/

DrugBank Database is available at the following URL: https://go.drugbank.com/gnomAD genic intolerance metrics are available at the following URL: (gnomAD v4.1.0) https://gnomad.broadinstitute.org/data

IEI-associated gene list 2024 is available at the following URL: https://iuis.org/committees/iei/

Druggable genome annotation is available at the following URL: doi.org/10.1126/scitranslmed.aag1166

Genetic Priority Score is available at the following URL: https://zenodo.org/records/10095684

HGNC complete gene set is available at the following URL: (accessed July 16, 2026) https://www.genenames.org/download/

STRING protein-protein interaction network is available at the following URL: https://string-db.org

Gene Ontology annotations are available at the following URL: http://geneontology.org

IGNITE ranked atlas for 19,502 genes is available (**Supplementary Table 1**).

## Code Availability

Code to construct the IGNITE score and to reproduce every figure and statistic reported here is available at https://github.com/alhanster/IGNITE_public. We performed all analyses in Python v3.12.13 using SciPy v1.18.0^87^, scikit-learn v1.9.0^88^ and XGBoost v3.3.0^89^ and generated figures with R v4.4.1 using ggplot2 v4.0.3^90^.

## Acknowledgments

We thank the participants and investigators who generated the Primary Human CD4⁺ T-Cell Perturb-seq resource. We also thank the gnomAD and Open Targets consortia. J.M.K. is supported by the National Institutes of Health through award NIAMS K24AR076975.

## Author Contributions

A.L.H. designed the study. A.L.H., J.W. performed analyses and statistical interpretation. A.L.H. performed data visualization. A.L.H. developed the web portal. A.L.H., M.S., D.A.F., R.S.D., J.W. wrote the manuscript. A.L.H., D.A.F., M.S., R.S.D., J.M.K., J.E.G., J.W. reviewed the manuscript.

## Ethics summary

This study complies with all relevant ethical regulations. It analyzes only publicly available, de-identified human data, and no ethical approval was required.

## References

1. Nelson, M. R. et al. The support of human genetic evidence for approved drug indications. Nat. Genet. 47, 856–860 (2015).

2. Ochoa, D. et al. Human genetics evidence supports two-thirds of the 2021 FDA-approved drugs. Nat. Rev. Drug Discov. 21, 551–551 (2022).

3. Razuvayevskaya, O., Lopez, I., Dunham, I. & Ochoa, D. Genetic factors associated with reasons for clinical trial stoppage. Nat. Genet. 56, 1862–1867 (2024).

4. King, E. A., Davis, J. W. & Degner, J. F. Are drug targets with genetic support twice as likely to be approved? Revised estimates of the impact of genetic support for drug mechanisms on the probability of drug approval. PLOS Genet. 15, e1008489 (2019).

5. Trajanoska, K. et al. From target discovery to clinical drug development with human genetics. Nature 620, 737–745 (2023).

6. Minikel, E. V., Painter, J. L., Dong, C. C. & Nelson, M. R. Refining the impact of genetic evidence on clinical success. Nature 629, 624–629 (2024).

7. Wang, Q. et al. Rare variant contribution to human disease in 281,104 UK Biobank exomes. Nature 597, 527–532 (2021).

8. Duffy, Á. et al. Development of a human genetics-guided priority score for 19,365 genes and 399 drug indications. Nat. Genet. 56, 51–59 (2024).

9. Chen, R. et al. Expanding drug targets for 112 chronic diseases using a machine learning-assisted genetic priority score. Nat. Commun. 15, 8891 (2024).

10. Fang, H. et al. A genetics-led approach defines the drug target landscape of 30 immune-related traits. Nat. Genet. 51, 1082–1091 (2019).

11. Fang, H. & Knight, J. C. Priority index: database of genetic targets in immune-mediated disease. Nucleic Acids Res. 50, D1358–D1367 (2022).

12. Weeks, E. M. et al. Leveraging polygenic enrichments of gene features to predict genes underlying complex traits and diseases. Nat. Genet. 55, 1267–1276 (2023).

13. Cornish, A. J., David, A. & Sternberg, M. J. E. PhenoRank: reducing study bias in gene prioritization through simulation. Bioinformatics 34, 2087–2095 (2018).

14. Eswaran, E., Zimmermann, M., Krishnagopalan, G., Ottersbach, J. M. & Gossmann, T. I. Evolutionary and functional constraints structure human gene research visibility. BMC Genomics 27, 526 (2026).

15. Balachandran, S. et al. STIGMA: Single-cell tissue-specific gene prioritization using machine learning. Am. J. Hum. Genet. 111, 338–349 (2024).

16. Chen, R., Duffy, Á. & Do, R. Genomics of drug target prioritization for complex diseases. Nat. Rev. Genet. 27, 231–245 (2026).

17. Falaguera, M. J. et al. Temporal trends in evidence supporting novel drug target discovery. Nat. Commun. 17, 492 (2025).

18. Alsentzer, E., Finlayson, S. G., Li, M. M., Kobren, S. N. & Kohane, I. S. Simulation of undiagnosed patients with novel genetic conditions. Nat. Commun. 14, 6403 (2023).

19. Freimer, J. W. et al. Systematic discovery and perturbation of regulatory genes in human T cells reveals the architecture of immune networks. Nat. Genet. 54, 1133–1144 (2022).

20. Kampmann, M. CRISPR-based functional genomics for neurological disease. Nat. Rev. Neurol. 16, 465–480 (2020).

21. GBD 2019 IMID Collaborators. Global, regional, and national incidence of six major immune-mediated inflammatory diseases: findings from the global burden of disease study 2019. EClinicalMedicine 64, 102193 (2023).

22. El-Gabalawy, H., Guenther, L. C. & Bernstein, C. N. Epidemiology of immune-mediated inflammatory diseases: incidence, prevalence, natural history, and comorbidities. J. Rheumatol. Suppl. 85, 2–10 (2010).

23. Kalden, J. R. & Schulze-Koops, H. Immunogenicity and loss of response to TNF inhibitors: implications for rheumatoid arthritis treatment. Nat. Rev. Rheumatol. 13, 707–718 (2017).

24. Pitzalis, C., Choy, E. H. S. & Buch, M. H. Transforming clinical trials in rheumatology: towards patient-centric precision medicine. Nat. Rev. Rheumatol. 16, 590–599 (2020).

25. Ding, N. S., Hart, A. & De Cruz, P. Systematic review: predicting and optimising response to anti-TNF therapy in Crohn’s disease - algorithm for practical management. Aliment. Pharmacol. Ther. 43, 30–51 (2016).

26. Veerasubramanian, P. K., Wynn, T. A., Quan, J. & Karlsson, F. J. Targeting TNF/TNFR superfamilies in immune-mediated inflammatory diseases. J. Exp. Med. 221, e20240806 (2024).

27. Gupta, A. et al. Dynamic regulatory elements in single-cell multimodal data implicate key immune cell states enriched for autoimmune disease heritability. Nat. Genet. 55, 2200–2210 (2023).

28. Soskic, B. et al. Chromatin activity at GWAS loci identifies T cell states driving complex immune diseases. Nat. Genet. 51, 1486–1493 (2019).

29. Mouri, K. et al. Prioritization of autoimmune disease-associated genetic variants that perturb regulatory element activity in T cells. Nat. Genet. 54, 603–612 (2022).

30. Harroud, A. & Hafler, D. A. Common genetic factors among autoimmune diseases. Science 380, 485–490 (2023).

31. Schmidt, R. et al. CRISPR activation and interference screens decode stimulation responses in primary human T cells. Science 375, eabj4008 (2022).

32. Zhu, R. et al. Genome-scale perturb-seq in primary human CD4+ T cells maps context-specific regulators of T cell programs and human immune traits. Cell 10.1016/j.cell.2026.08.002 (2026) doi:10.1016/j.cell.2026.08.002.

33. Morris, J. A. et al. Discovery of target genes and pathways at GWAS loci by pooled single-cell CRISPR screens. Science 380, eadh7699 (2023).

34. Augustine, M. et al. Immunotherapy drug target identification using machine learning and patient-derived tumour explant validation. Nat. Mach. Intell. 8, 670–689 (2026).

35. Ota, M. et al. Causal modelling of gene effects from regulators to programs to traits. Nature 650, 399–408 (2026).

36. Li, F. et al. Positive-unlabeled learning in bioinformatics and computational biology: a brief review. Brief. Bioinform. 23, bbab461 (2022).

37. Muslu, O., Hoyt, C. T., Lacerda, M., Hofmann-Apitius, M. & Frohlich, H. GuiltyTargets: Prioritization of Novel Therapeutic Targets With Network Representation Learning. IEEE/ACM Trans. Comput. Biol. Bioinform. 19, 491–500 (2022).

38. Bousfiha, A. A. et al. The 2024 update of IUIS phenotypic classification of human inborn errors of immunity. J. Hum. Immun. 1, e20250002 (2025).

39. Buniello, A. et al. Open Targets Platform: facilitating therapeutic hypotheses building in drug discovery. Nucleic Acids Res. 53, D1467–D1475 (2025).

40. Integrating 730,947 exome sequences with clinical literature improves gene discovery | medRxiv. https://www.medrxiv.org/content/10.64898/2026.03.23.26349081v1.

41. Finan, C. et al. The druggable genome and support for target identification and validation in drug development. Sci. Transl. Med. 9, eaag1166 (2017).

42. Pencina, M. J., D’Agostino, R. B., Pencina, K. M., Janssens, A. C. J. W. & Greenland, P. Interpreting incremental value of markers added to risk prediction models. Am. J. Epidemiol. 176, 473–481 (2012).

43. Sadler, M. C., Auwerx, C., Deelen, P. & Kutalik, Z. Multi-layered genetic approaches to identify approved drug targets. Cell Genomics 3, 100341 (2023).

44. Yang, P., Li, X.-L., Mei, J.-P., Kwoh, C.-K. & Ng, S.-K. Positive-unlabeled learning for disease gene identification. Bioinformatics 28, 2640–2647 (2012).

45. Hu, Y., Zhao, T., Zhang, N., Zhang, Y. & Cheng, L. A Review of Recent Advances and Research on Drug Target Identification Methods. Curr. Drug Metab. 20, 209–216 (2019).

46. Kolosov, N., Daly, M. J. & Artomov, M. Prioritization of disease genes from GWAS using ensemble-based positive-unlabeled learning. Eur. J. Hum. Genet. 29, 1527–1535 (2021).

47. Pahikkala, T. et al. Toward more realistic drug-target interaction predictions. Brief. Bioinform. 16, 325–337 (2015).

48. Zhang, S., Wang, J., Lin, Z. & Liang, Y. Application of Machine Learning Techniques in Drug-target Interactions Prediction. Curr. Pharm. Des. 27, 2076–2087 (2021).

49. Knox, C. et al. DrugBank 6.0: the DrugBank Knowledgebase for 2024. Nucleic Acids Res. 52, D1265–D1275 (2024).

50. Baghdassarian, H. et al. Variant STAT4 and Response to Ruxolitinib in an Autoinflammatory Syndrome. N. Engl. J. Med. 388, 2241–2252 (2023).

51. Brovchenko, N. et al. Biaryl Phosphates and Phosphonates as Selective Inhibitors of the Transcription Factor STAT4. Angew. Chem. 64, e202504420 (2025).

52. Remmers, E. F. et al. STAT4 and the risk of rheumatoid arthritis and systemic lupus erythematosus. N. Engl. J. Med. 357, 977–986 (2007).

53. O’Shea, J. J. et al. The JAK-STAT pathway: impact on human disease and therapeutic intervention. Annu. Rev. Med. 66, 311–328 (2015).

54. Miklossy, G., Hilliard, T. S. & Turkson, J. Therapeutic modulators of STAT signalling for human diseases. Nat. Rev. Drug Discov. 12, 611–629 (2013).

55. Szalai, T. V. et al. Allosteric Covalent Inhibitors of the STAT3 Transcription Factor from Virtual Screening. ACS Med. Chem. Lett. 16, 991–997 (2025).

56. Jung, H. & Lee, Y. Targeting the Undruggable: Recent Progress in PROTAC-Induced Transcription Factor Degradation. Cancers 17, 1871 (2025).

57. Szeląg, M., Czerwoniec, A., Wesoly, J. & Bluyssen, H. A. R. Comparative screening and validation as a novel tool to identify STAT-specific inhibitors. Eur. J. Pharmacol. 740, 417–420 (2014).

58. Furtek, S. L., Backos, D. S., Matheson, C. J. & Reigan, P. Strategies and Approaches of Targeting STAT3 for Cancer Treatment. ACS Chem. Biol. 11, 308–318 (2016).

59. Attarha, S., Reithmeier, A., Busker, S., Desroses, M. & Page, B. D. G. Validating Signal Transducer and Activator of Transcription (STAT) Protein-Inhibitor Interactions Using Biochemical and Cellular Thermal Shift Assays. ACS Chem. Biol. 15, 1842–1851 (2020).

60. Tsimberidou, A. M. et al. Phase I Trial of TTI-101, a First-in-Class Oral Inhibitor of STAT3, in Patients with Advanced Solid Tumors. Clin. Cancer Res. Off. J. Am. Assoc. Cancer Res. 31, 965–974 (2025).

61. Shimamura, K. et al. 5,5-Dimethyl-3-(5-methyl-3-oxo-2-phenyl-2,3-dihydro-1H-pyrazol-4-yl)-1-phenyl-3-(trifluoromethyl)-3,5,6,7-tetrahydro-1H-indole-2,4-dione, a potent inhibitor for mammalian elongase of long-chain fatty acids family 6: examination of its potential utility as a pharmacological tool. J. Pharmacol. Exp. Ther. 330, 249–256 (2009).

62. Bantug, G. R., Galluzzi, L., Kroemer, G. & Hess, C. The spectrum of T cell metabolism in health and disease. Nat. Rev. Immunol. 18, 19–34 (2018).

63. Kanno, T., Nakajima, T., Miyako, K. & Endo, Y. Lipid metabolism in Th17 cell function. Pharmacol. Ther. 245, 108411 (2023).

64. Sharabi, A. & Tsokos, G. C. T cell metabolism: new insights in systemic lupus erythematosus pathogenesis and therapy. Nat. Rev. Rheumatol. 16, 100–112 (2020).

65. Batchuluun, B., Pinkosky, S. L. & Steinberg, G. R. Lipogenesis inhibitors: therapeutic opportunities and challenges. Nat. Rev. Drug Discov. 21, 283–305 (2022).

66. Yoshida, K. et al. ELOVL6 deficiency aggravates allergic airway inflammation through the ceramide-S1P pathway in mice. J. Allergy Clin. Immunol. 151, 1067–1080.e9 (2023).

67. Kiyoki, Y. et al. The fatty acid elongase Elovl6 is crucial for hematopoietic stem cell engraftment and leukemia propagation. Leukemia 37, 910–913 (2023).

68. Garcia Corrales, A. V., et al. Fatty acid elongation by ELOVL6 hampers remyelination by promoting inflammatory foam cell formation during demyelination. Proc. Natl. Acad. Sci. U. S. A. 120, e2301030120 (2023).

69. Matsuzaka, T. et al. Elovl6 promotes nonalcoholic steatohepatitis. Hepatology 56, 2199–2208 (2012).

70. Shimamura, K. et al. Discovery and characterization of a novel potent, selective and orally active inhibitor for mammalian ELOVL6. Eur. J. Pharmacol. 630, 34–41 (2010).

71. Nagase, T. et al. Synthesis and biological evaluation of a novel 3-sulfonyl-8-azabicyclo[3.2.1]octane class of long chain fatty acid elongase 6 (ELOVL6) inhibitors. J. Med. Chem. 52, 4111–4114 (2009).

72. Mizutani, T. et al. Discovery of novel benzoxazinones as potent and orally active long chain fatty acid elongase 6 inhibitors. J. Med. Chem. 52, 7289–7300 (2009).

73. Takahashi, T. et al. Synthesis and evaluation of a novel indoledione class of long chain fatty acid elongase 6 (ELOVL6) inhibitors. J. Med. Chem. 52, 3142–3145 (2009).

74. Sasaki, T. et al. Synthesis and evaluation of a novel 2-azabicyclo[2.2.2]octane class of long chain fatty acid elongase 6 (ELOVL6) inhibitors. Bioorg. Med. Chem. 17, 5639–5647 (2009).

75. Matias, P. M., Gorynia, S., Donner, P. & Carrondo, M. A. Crystal structure of the human AAA+ protein RuvBL1. J. Biol. Chem. 281, 38918–38929 (2006).

76. Dauden, M. I., López-Perrote, A. & Llorca, O. RUVBL1-RUVBL2 AAA-ATPase: a versatile scaffold for multiple complexes and functions. Curr. Opin. Struct. Biol. 67, 78–85 (2021).

77. Gnatovskiy, L., Mita, P. & Levy, D. E. The human RVB complex is required for efficient transcription of type I interferon-stimulated genes. Mol. Cell. Biol. 33, 3817–3825 (2013).

78. Ivashkiv, L. B. & Donlin, L. T. Regulation of type I interferon responses. Nat. Rev. Immunol. 14, 36–49 (2014).

79. Kaji, K. et al. Autoantibodies to RuvBL1 and RuvBL2: a novel systemic sclerosis-related antibody associated with diffuse cutaneous and skeletal muscle involvement. Arthritis Care Res. 66, 575–584 (2014).

80. Cavazzana, I. et al. Systemic Sclerosis-Specific Antibodies: Novel and Classical Biomarkers. Clin. Rev. Allergy Immunol. 64, 412–430 (2023).

81. Assimon, V. A. et al. CB-6644 Is a Selective Inhibitor of the RUVBL1/2 Complex with Anticancer Activity. ACS Chem. Biol. 14, 236–244 (2019).

82. Vogt, M. et al. Targeting MYC effector functions in pancreatic cancer by inhibiting the ATPase RUVBL1/2. Gut 73, 1509–1528 (2024).

83. Minikel, E. V. et al. Evaluating drug targets through human loss-of-function genetic variation. Nature 581, 459–464 (2020).

84. Spradlin, J. N., Zhang, E. & Nomura, D. K. Reimagining Druggability Using Chemoproteomic Platforms. Acc. Chem. Res. 54, 1801–1813 (2021).

85. Sun, Q. et al. Computer-Aided Drug Discovery for Undruggable Targets. Chem. Rev. 125, 6309–6365 (2025).

86. Han, A. L. et al. Diverse ancestral representation improves genetic intolerance metrics. Nat. Commun. 16, 2648 (2025).

87. Virtanen, P. et al. SciPy 1.0: fundamental algorithms for scientific computing in Python. Nat. Methods 17, 261–272 (2020).

88. Pedregosa, F., et al. Scikit-learn: Machine Learning in Python. Preprint at 10.48550/arXiv.1201.0490 (2018).

89. Chen, T. & Guestrin, C. XGBoost: A Scalable Tree Boosting System. in Proceedings of the 22nd ACM SIGKDD International Conference on Knowledge Discovery and Data Mining 785–794 (2016). doi:10.1145/2939672.2939785.

90. ggplot2: Elegant Graphics for Data Analysis (3e). https://ggplot2-book.org/.

