## Supplementary Information for "Genome-scale perturbation signatures from primary human CD4^+^ T cells improve genetics-based prioritization of immune drug targets"

### Supplementary Figure 1

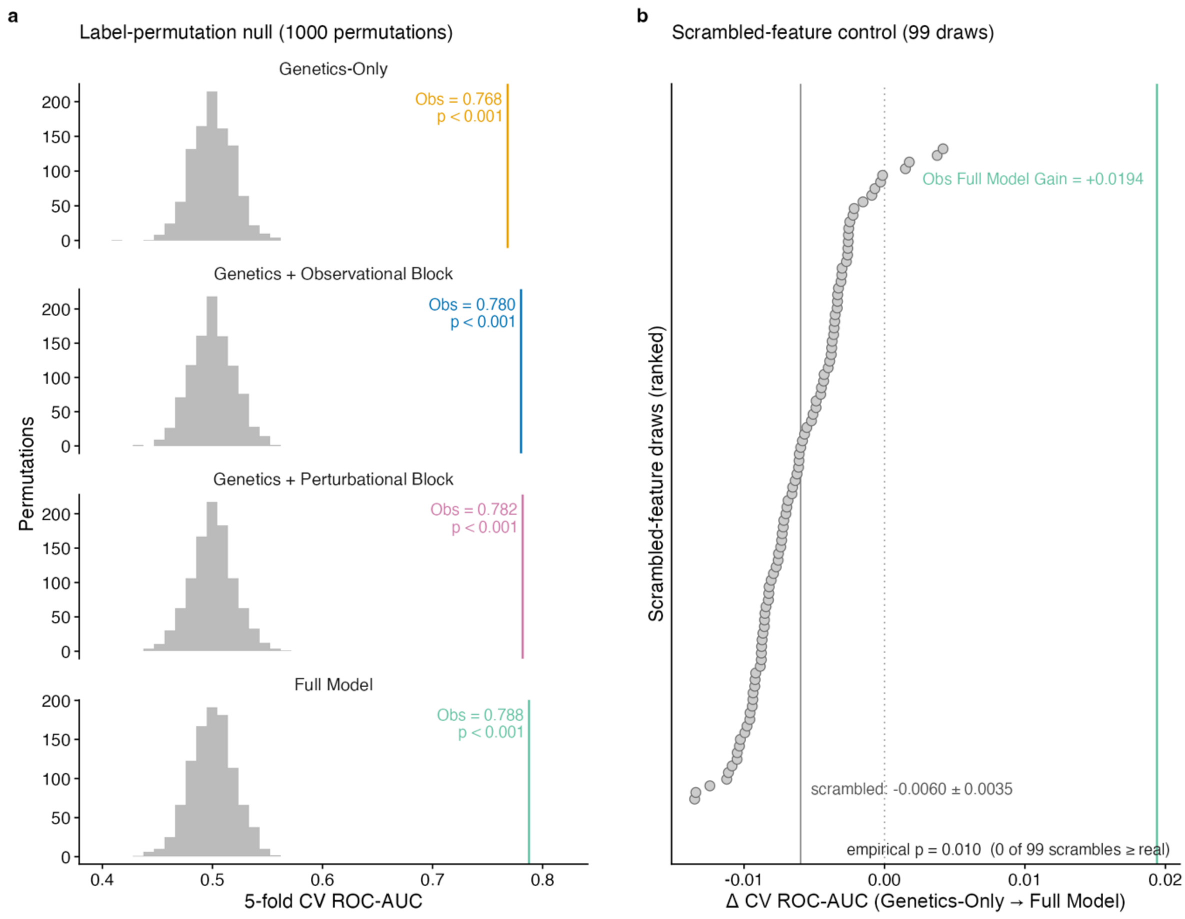

**Label-permutation and scrambled-feature nulls. (a)** Label-permutation null over 1,000 permutations for each evidence rung (genetics only, genetics + observational functional genomics, genetics + perturbational functional genomics, full model). The 340 positives are reassigned at random, and the full cross-validation pipeline is rerun for each rung. Null distributions center on AUC 0.500 with standard deviations 0.019-0.020, and each rung sits at least 14 standard deviations above the null (empirical $p < 0.001$ at all four rungs). This panel calibrates the coalition levels. **(b)** Scrambled-feature control for the functional-genomic increment, the appropriate null for the increment. True labels and true genetic features are held fixed while each of 19 functional-genomic features is independently permuted across rows (99 draws). The observed increment of +0.0194 AUC exceeded all 99 scrambled draws (mean -0.0060 ± 0.0035, empirical $p = 0.010$).

### Supplementary Figure 2

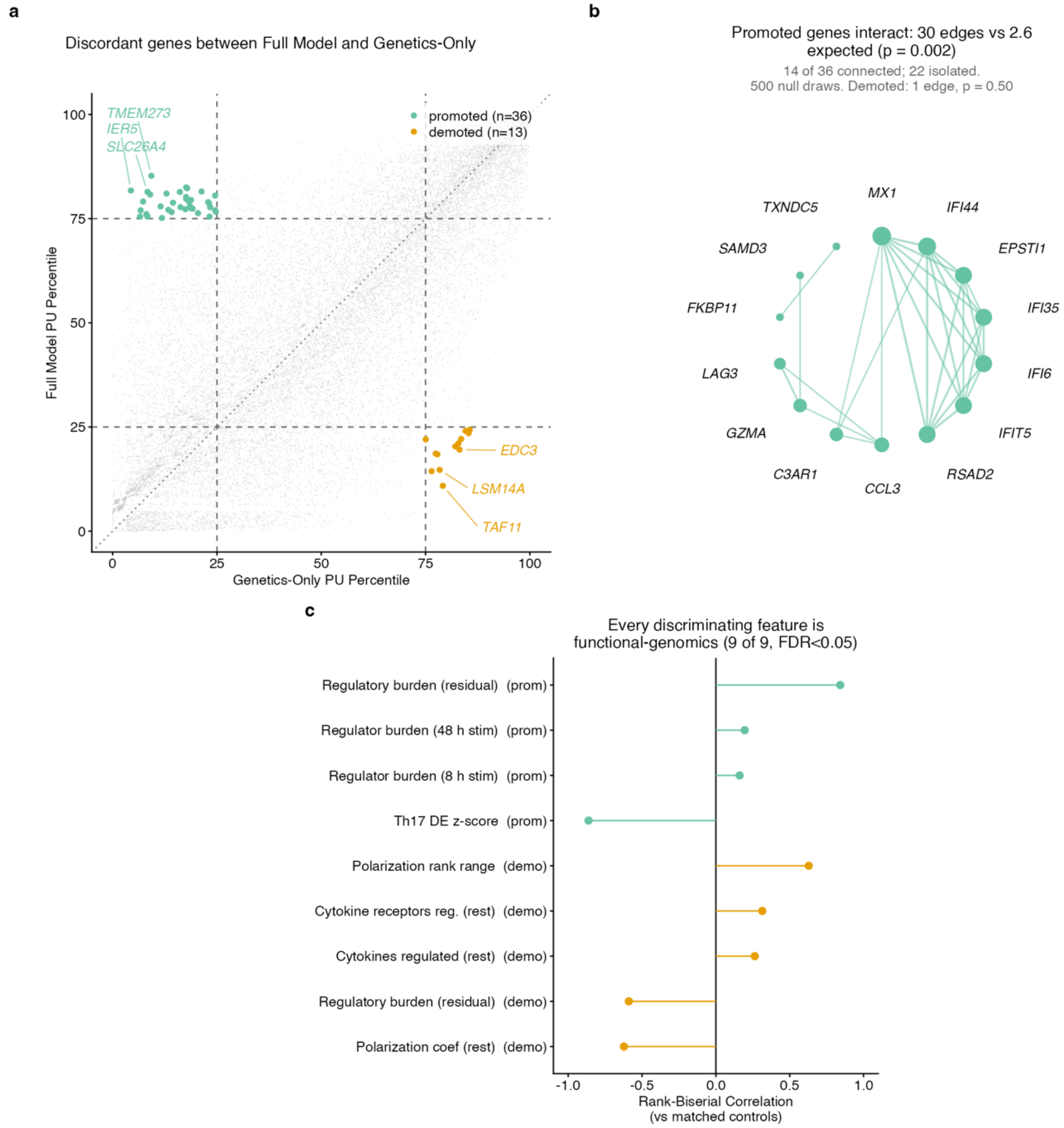

**Genes ranked discordantly by the genetics-only and full models. (a)** Rank percentile under the full model versus the genetics-only model for all 19,502 genes (Spearman $\rho = 0.83$). The 25th and 75th percentile boundaries are drawn and the 49 discordant genes are marked. **(b)** STRING protein–protein interaction edges among the promoted genes against 500 matched null gene sets. The demoted set shows no such structure. **(c)** Features distinguishing promoted or demoted genes from pooled matched controls, as rank-biserial effect sizes. All nine discriminating features (FDR < 0.05) are functional-genomic. The largest promoted-direction effect is Th17 differential expression, followed by residualized trans-regulatory burden. Residualized trans-regulatory burden is the only feature that separates in both directions.

### Supplementary Figure 3

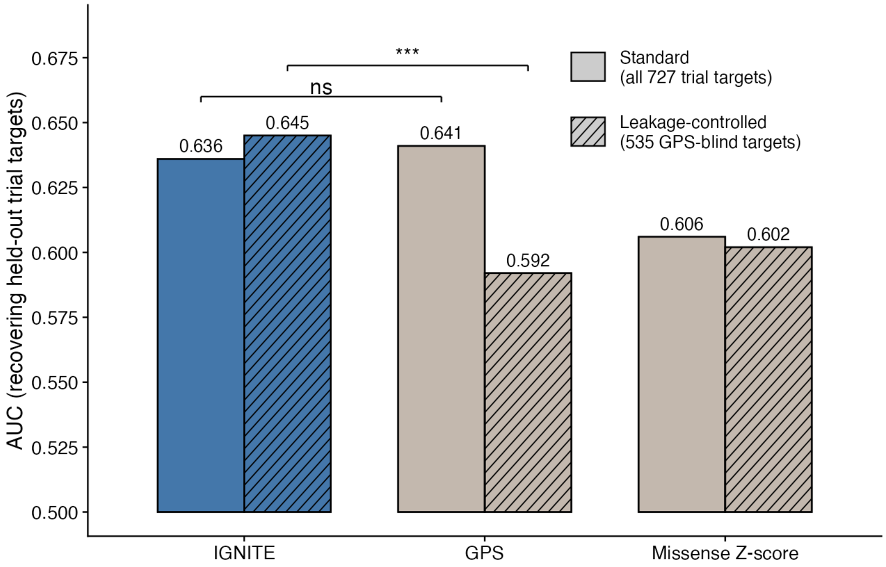

**Standard and leakage-controlled comparison against published genetic priority scores.** Recovery of held-out immune-trial targets by IGNITE, maximum per-gene Genetic Priority Score (GPS), and gnomAD missense Z-score. The standard arm scores all 727 held-out in-trial genes, ranked by IGNITE and GPS across 19,162 genes and by missense-Z across 19,087. The two counts differ because a gene with no published GPS is scored zero while a gene with no missense Z-score is dropped. IGNITE and GPS are statistically indistinguishable on this arm by a paired DeLong test. The leakage-controlled arm removes the 284 genes GPS used as positive drug labels in training that fall inside our non-approved pool, leaving 535 in-trial genes across 18,878 genes (18,803 for missense-Z) that neither method trained on as positives. For the leakage-controlled set, IGNITE reaches 0.645 against 0.592 for GPS which was significant based on paired DeLong test. Missense Z-score is shown as an unsupervised constraint reference and is unaffected by the leakage control. Brackets compare IGNITE with GPS within each arm. Bars start at 0.5 (chance), not at zero. * $p < 0.05$, ** $p < 0.01$, *** $p < 10^{-3}$, **** $p < 10^{-4}$; ns, not significant.

### Supplementary Table Legends

#### Supplementary Table 1 – IGNITE ranked atlas for 19,502 genes

Provided as external file. Genes and their IGNITE positive-unlabeled (PU) prioritization scores are described in the table, one row per gene. Variable "pu_score" is the out-of-bag PU prioritization score (Methods). Values for the 340 approved genes are in-bag and are reported only so the ranking is complete. Variables "rank" and "rank_pctile" give the genome-wide rank (1 = highest score) and rank percentile. Variables "drug_status" and "furthest_imm_stage" are as defined in Supplementary Table 3, and variable "pu_role" gives the role of the gene in model fitting (P = training positive, trial_heldout = held-out in-trial, unlabeled). Columns for the 24 modeled features follow the names, definitions, evidence-block assignment, and stimulation condition given in Supplementary Table 2. Abbreviations used in column names and in Supplementary Table 2: "lof.oe_ci.upper" = LOEUF = loss-of-function observed/expected upper bound fraction. Variables "crossdonor_confidence" and "crossdonor_correlation_mean" carry the cross-donor reproducibility annotation of the perturbation signature and are identical by construction.

#### Supplementary Table 2 – Feature dictionary and evidence-block membership

Provided as external file. The 24 modeled features are described in the table, one row per feature. Variable "feature" is the column name as it appears in the feature matrix and in Supplementary Table 1, and variable "block" assigns the feature to one of three evidence blocks (genetic, 5 features; observational, 4 features; perturbational, 15 features). Variable "condition" indicates the stimulation context of the measurement (Rest, Stim8hr, Stim48hr, or one of three across-condition summaries: coverage, mean of 3, range of 3) and is empty for features without a condition. Variable "type" records whether the feature is binary or continuous, variable "definition" gives the quantity measured and any transformation applied, and variable "observed_rho" the observed Spearman correlation between the feature and the full model score over genes with an observed value. Variables "n_nonmissing" and "pct_nonmissing" account for the number and percentage of the 19,502 genes with an observed value. Variable "imputation" states the rule applied to missing entries, and variable "source" names the originating resource and release. Variables "imp_mean" and "imp_sd" are the per-feature normalized gain importance across bagging iterations and its population standard deviation.

#### Supplementary Table 3 – Drug-status labels and immune-indication trial annotations

Provided as external file. Genes and their immune-indication development status are described in the table, one row per gene with any immune-indication drug annotation (1,089 genes). Variable "drug_status" is the assigned label (approved, in-trial, non-target) and variable "furthest_imm_stage" the most advanced immune-indication phase reached by any drug annotated to the gene. Variable "n_drugs" accounts for the total number of immune-indication drugs annotated to the gene, variable "drug_names" lists them, and variable "indications" lists the corresponding disease terms. Variable "first_trial_year" gives the earliest trial start year across those drugs in any indication (not just immune indication) and variable "approval_year" the year of first approval for immune-indication (for 332 of the 340 approved genes). Variable "multi_target_expanded" flags genes entered through expansion of a multi-target drug and variable "in_feature_matrix" indicates presence in the 19,502-gene feature matrix. Labels and trial annotations were derived from the Open Targets Platform 26.06 evidence tables.

#### Supplementary Table 4 – Univariate feature statistics by drug status

Provided as external files. Each panel of Figures 2 and 3 is present on separate sheets, accompanied by test and summary sheets. Sheets from left to right: "Fig 2a IEI enrichment", "Fig 2b immune-GWAS", "Fig 2b pct above zero", "Fig 2b tests", "Fig 2c missense Z", "Fig 2c tests", "Fig 3a T-helper DE", "Fig 3a FDR enrichment", "Fig 3b regulatory residual", "Fig 3b residual tests", "Fig 3c cytokines", "Fig 3d cytokine receptors", "Fig 3cd distributions", "Fig 3cd regulator fraction", and "Group sizes".

On the gene-level sheets, variable "group" is the drug-status group (approved, in-trial, non-target) and variables "gwas_score", "mis_z_score", "neglog10_adjp", "residual" and "n_sig" carry the immune-GWAS (genome-wide association study) score, the missense Z-score, the negative base-10 logarithm of the adjusted differential-expression p-value, the trans-regulatory burden residual and the count of significantly regulated cytokines or cytokine receptors, respectively. The immune-GWAS sheet covers all 19,502 genes. The missense Z sheet covers 19,427 genes. Variable "subset" on the T-helper sheets is the polarized subset compared against resting Th0.

On the enrichment and test sheets, variable "background" names the comparison set on the IEI enrichment sheet. Variables "n", "n_iei", "n_tested", "n_hit" and "pct_hit" give group sizes and hit counts. Variables "n_gt0_a", "n_gt0_b", "pct_gt0_a", "pct_gt0_b" and "pct_gt0" indicate the counts and percentages scoring above zero. Variables "OR", "OR_lo" and "OR_hi" provide the Fisher sample odds ratio with its Woolf 95% confidence interval (CI). Variables "median_a", "median_b", "hl_shift", "hl_lo" and "hl_hi" indicate the group medians and the Hodges-Lehmann location shift with its 95% interval. Variables "p_sup", "p_sup_lo" and "p_sup_hi" are the probability of superiority with its Hanley-McNeil 95% interval. Variables "statistic" and "value" are the test statistic and its value. Variables "p", "p_holm" and "stars" are the raw p-value, the Holm-Bonferroni corrected p-value and the corresponding star rung, respectively. Variable "genes" on the Figure 2b test sheet records which gene group each row was computed on. This sheet contains two separately corrected blocks: three Fisher tests on all genes, asking which genes carry any immune-GWAS signal, and a Kruskal-Wallis omnibus with three pairwise Mann-Whitney tests on the genes scoring above zero. The Figure 2c and 3b test sheets lead with the same omnibus row; omnibus rows are uncorrected and carry empty pairwise cells.

On the Figure 3c/3d summary sheets, variables "n_reg", "pct_reg" and "pct_zero" describe how often a gene regulates at least one target. Variables "mean" and "mean_nonzero" indicate the group mean overall and among regulating genes. Variables "max" and "n_ge3" report the largest count and the number of genes regulating three or more targets. Variables "n_ref", "n_reg_ref" and "pct_reg_ref" on the regulator-fraction sheet describe the non-target genes each class is tested against in the same condition. Group sizes for every panel are given in the final sheet by variables "panel", "group" and "n". Tests and their correction families are described in Methods.

#### Supplementary Table 5 – Orthogonality of functional-genomic evidence to genetic priors

Provided as external files. Each analysis is present on separate sheets. "Genetic vs functional genomic" sheet describes every pairing of a genetic prior with a functional-genomic feature. Variable "genetic" names the genetic prior, and the variable "fg" names the functional-genomic feature. Variable "n" accounts for the number of genes with both values observed. Variable "rho" is the Spearman correlation coefficient with its p-value in variable "p". Variable "shared_var_pct" is the percentage of shared variance implied by that coefficient. "Among genetic priors" sheet describes the 10 pairings among the five genetic priors themselves: variables "feature_1" and "feature_2" name the pair. Variables "axis_1" and "axis_2" assign each to an evidence axis (rare-variant (curated), rare-variant (burden), common-variant, or evolutionary constraint). Variable "relation" records whether the pair spans two axes or sits within one, and variables "spearman_rho", "spearman_p", "pearson_r" and "shared_variance_pct_spearman" report the correlation statistics. These correlations are descriptive and are reported uncorrected.

**Supplementary Table 6 – Cross-validated performance of all evidence coalitions and attribution**

Provided as external files. Each analysis is present on separate sheets. “Coalitions” sheet describes all eight coalitions of the three evidence blocks, one row per coalition. Variable "coalition" names the blocks included, and variables "auc_mean" and "auc_sd" are the mean cross-validated AUC (area under the receiver operating characteristic curve) and its population standard deviation across the 25 samples. In the “Model ladder” sheet, variable "model" identifies the model variant, variable "label" its display label, and variables "auc", "sd" and "sem" the mean AUC, standard deviation and standard error. The “Significance” sheet reports variables "model_a" and "model_b" for the compared variants, variable "delta" for the paired change in AUC, variable "nb_p" for the Nadeau-Bengio corrected p-value, and variable "stars" for the corresponding annotation. Variables in “Attribution” sheet are "evidence" (evidence block), "shapley_auc" (grouped-Shapley value of AUC above chance) and "share_pct" (its percentage share). In the “Perturbational sub-block strata” sheet, variables "weak_lift" and "strong_lift" are the incremental signal contributed within the weak- and strong-genetic-prior strata. Variable "block" names the six perturbational feature groups rather than the three evidence blocks of the other sheets. Model configurations, the Shapley decomposition and the Nadeau-Bengio test are described in Methods.

**Supplementary Table 7 – Recovery of held-out in-trial targets and score distribution by drug status**

Provided as external files. Each analysis is present on separate sheets. In the “Recovery at depth” sheet, variable "k" is the nomination depth with variable "depth" its display label. Variables "genetics" and "genetics_perturbseq" account for the number of the 727 held-out in-trial genes recovered by the genetics-only and full models, and variable "gain" reports their difference. The “ROC curves” sheet gives variables "model", "fpr" and "tpr". The AUCs, their DeLong confidence intervals and the paired DeLong test are computed on the full curves and reported in the “Discrimination statistics” sheet by variables "statistic", "model", "value", "ci_lo" and "ci_hi". The "Score distribution" sheet describes all 19,502 genes by variables "gene", "grp" (drug-status group). Variables "pu_score" and "rank_pctile" are the published model score and rank percentile. Variables "pu_score_cv" and "rank_pctile_cv" are the cross-fitted score and rank percentile where every gene, including the 340 approved training positives, is scored by models that did not train on it using five folds over the approved genes. The "Group summary" sheet reports each group's size and the median, mean and quartiles of "rank_pctile_cv" by variables "group", "n", "median", "mean", "q25" and "q75". The "Group tests" sheet reports, on "pu_score_cv", the Kruskal-Wallis omnibus test, the one-way ANOVA, the one-sided Mann-Whitney test of held-out in-trial against non-target genes, and the three pairwise Mann-Whitney tests, by variables "test", "group_a", "group_b", "statistic", "value", "p", "p_holm" and "stars". Only the three pairwise tests are corrected, so variables "p_holm" and "stars" are empty on the other rows.

**Supplementary Table 8 – Label-permutation null and scrambled-feature control**

Provided as external files. Each analysis is present on separate sheets. The “Permutation null” sheet describes the 1,000 label permutations, one row per permutation. Variable "perm" is the permutation index, and variables "genetic", "genetic+observational", "genetic+perturbational" and "full" give the AUC obtained by each model variant under that reassignment of positives. Variables "obs_over_genetic", "perturbational_over_genetic" and "full_over_genetic" describe the corresponding increments over the genetics-only variant. The “Summary” sheet describes each tested quantity. Variable "metric" and “observed” indicate the model variant and its observed value. Variables "null_mean", "null_sd" and "null_p95" describe the null distribution. Variable "emp_p" is the empirical p-value, and variable "n_perm" the number of permutations drawn. The “Scrambled-feature control” sheet reports its 99 draws by variable "kind", separating the observed increment from the individual draws and the summary. Variable "key" names the draw index or the summary quantity, and variable "value" reports the increment in AUC. Its empirical p-value of 0.010 is the floor attainable at that number of draws. Both nulls are described in Methods.

#### Supplementary Table 9 – Genes ranked discordantly by the genetics-only and full models

Provided as external files. Each analysis is present on separate sheets. Sheets from left to right: “Rank shift scatter”, “Core discordant genes”, “Feature tests”, “Rank-shift drivers”, “STRING network test”, “STRING edges”, “STRING nodes”, “GO enrichment”, “Annotation coverage”, “Annotation coverage axes”, and “Stratified validation”. The “Rank shift scatter” sheet describes all 19,502 genes where variables "pctile_genetics" and "pctile_full" are the rank percentiles under the two models. Variable "rank_shift" reports their difference. Variable "core" is the discordance label (promoted, 36 genes; demoted, 13; none, 19,453). Variable "gen_dec" is the genetic-prior decile, and variable "cov_class" is the annotation-coverage class. The "Core discordant genes" sheet describes these 49 genes by variables "gene", "core", "pctile_genetics", "pctile_full" and "rank_shift", their genetic and functional-genomic values ("IEI", "gwas_score", "mis.z_score", "lof.oe_ci.upper", "expected_n_regulators_residuals") and status flags ("is_druggable", "is_approved", "is_trial_heldout"). The “Feature tests” sheet describes each tested feature by variables "direction" (promoted or demoted), "block", "n_sel", "med_sel" and "med_ctrl" (group size and median in the selected and matched-control sets), "rank_biserial" (effect size), "p" and "fdr" (false discovery rate). The “Rank-shift drivers” sheet reports variables "rho", "p" and "n_nonnull" for the correlation of each feature with rank shift. The “STRING (Search Tool for the Retrieval of Interacting Genes/Proteins) network test” sheet describe the protein-protein interaction test by variables "direction", "n_genes", "n_mapped", "n_edges", "n_connected", "null_mean_edges", "null_max_edges", "fold", "p_empirical" and "n_null_draws". The “STRING edges” and “STRING nodes” sheets report the edge and node listings in variables "a", "b", "score" and "degree". The "GO enrichment" sheet (GO, Gene Ontology) reports variables "direction", "go_id", "term", "k_fg", "n_fg", "k_bg", "n_bg", "odds_ratio", "p", "fdr" and "n_family", where the "fg" and "bg" suffixes denote the foreground and background gene sets. The “Annotation coverage” sheets report variables "n_dark" and "pct_dark" for genes without a GO biological-process term in each of five groups, with variables "OR_vs_promoted" and "fisher_p" comparing the promoted set against each of the four reference groups and empty on the promoted row itself. Variables "axis", "comparison", "odds_ratio", "p", "verdict" and "background" are reported on the accompanying “Annotation coverage axes” sheet. The “Stratified validation” sheet reports variables "gen_stratum", "n", "pct_up", "pct_down", "k_up", "k_down", "odds_ratio" and "fisher_p" per genetic-prior stratum. The discordance definition, the matched controls and the two GO backgrounds are described in Methods.

#### Supplementary Table 10 – Top 50 IGNITE nominations and bootstrap trial enrichment

Provided as external files. Sheets from left to right: “Fig 5a nomination stages”, “Fig 5b top-50 enrichment”, and “Fig 5c enrichment by depth”. The “Fig 5a nomination stages” sheet describes the top 50 nominations excluding already approved immune targets, one row per nomination. Variable "rank_nonapproved" is the nomination rank after removal of approved targets. Variables "pu_score", "rank" and "rank_pctile" report the model score with its genome-wide rank and percentile. Variable "pu_role" indicates the gene's role in model fitting, and variable "furthest_imm_stage" is the furthest immune-indication trial stage if present. Variable "crossdonor_confidence" describes the cross-donor reproducibility annotation of the perturbation signature, with the "donor_vetted" variable recording greater than 0.3 as “yes”, lower as “weak”, and “not_measured” if absent. The “Fig 5b top-50 enrichment” sheet describes the bootstrap enrichment of the top 50 nominations of the full model and genetics-only model at two trial-stage "threshold" (Phase 1 or later, Phase 3 or later). Variables "observed_count" and "expected_count" report the observed and expected numbers of the 50 with such a drug. Variables "observed_pct" and "expected_pct" are the same quantities as percentages. Variable "fold_enrichment" describes their ratio, and variables "ci95_lo_pct" and "ci95_hi_pct" indicate the 2.5-97.5% bootstrap interval. Variable "p_bootstrap" is the empirical proportion of draws reaching the observed rate, and "p_bootstrap_bh" is proportion after Benjamini-Hochberg correction. Variables "n_top", "n_ctrl_per_draw", "n_bootstrap", "ctrl_pool_rank_min", "ctrl_pool_n" and "seed" record the sampling settings described in Methods. The “Fig 5c enrichment by depth” sheet repeats the any-immune-trial arm across shortlist depth. There is one row per combination of variable "depth" (25, 50 or 100) and variable "model".

#### Supplementary Table 11 – Comparison against Genetic Priority Scores with and without leakage control

Provided as external files. Sheets from left to right: “Comparison”, “DeLong statistics”, and “GPS drug labels”. The “Comparison” sheet describes each model score by evaluation combination. Variable "method" names the score where "PU full model" is IGNITE, "GPS overall" is the maximum per-gene Genetic Priority Score (GPS), and "missense-z" is the gnomAD missense Z-score as a single-feature comparator. Variable "evaluation" is the test set where "standard_taskA" is for all 727 held-out in-trial genes, and "leakage_controlled" is for the 535 in-trial genes unseen as positives by either method. Variable "auc" is the AUC. Variables "n_genes" and "n_pos" are the size of the evaluated universe and of the positive set. Variable "missing_policy" describes the handling of genes without a score, and variable "note" reports any qualification applying to that row. Variable "n_genes" is not constant in the sheet because the missing policy differs by method. A missing GPS score is scored as zero, while a missing missense Z-score is dropped. Paired DeLong statistics for the contrast of IGNITE against GPS are reported in the “DeLong statistics” sheet by variables "arm", "statistic" and "value". One block per evaluation arm, reporting the two AUCs, their difference, z, the raw p-value, and p-value after Holm-Bonferroni correction. The “GPS drug labels” sheet lists the 453 genes used as positive drug labels in GPS training, which are removed to define the leakage-controlled arm. The leakage-control procedure is described in Methods. GPS values were derived from Duffy et al., 2024.

#### Supplementary Table 12 – Prospective temporal holdout composition and results

Provided as external files. Sheets from left to right: “Scores at T0”, “Emergent genes”, “Headline AUC”, and “Freeze-year sensitivity”. The “Scores at T0” sheet describes all 19,502 genes at the headline freeze year *T0* = 2014. Variables "score_full", "rank_pctile_full", "score_genetics" and "rank_pctile_genetics" give the scores and rank percentiles of the full and genetics-only models retrained on pre-*T0* labels. Variables "is_pos_at_T0", "is_emergent", "is_holdout" and "in_pool" indicate whether the gene was a training positive at T0, an emergent gene, an excluded labeled target, or a member of the unlabeled evaluation pool, respectively. The “Emergent genes” sheet describes the 134 emergent genes at *T0* = 2014 by variable "first_trial_year" and the two rank percentiles repeated from the “Scores at T0” sheet. The “Headline AUC” sheet gives each model's emergent-vs-pool AUC at *T0* = 2014 in variable "auc", with the permutation-null p-value in variable "p_emp" and the emergent count in variable "n_emergent". The “Freeze-year sensitivity” sheet describes freeze years 2013 to 2017, one row per year. Variable "T0" is the freeze year, and variable "n_P" accounts for the number of training positives. Variable "n_emergent" reports the number of emergent genes at *T0*. Variables "auc_full" and "auc_genetics" give each model's AUC. Variable "delta" indicates their difference with its standard error in "se_delta". Variables "delong_p_two_sided" and "delong_p_one_sided" are the paired DeLong p-values. Variables "ci95_lo" and "ci95_hi" are the 95% CI of the AUC difference. Variable "p_full" is the permutation-null p-value for the full model. The freeze-year design is described in Methods. Trial dates were derived from the Open Targets Platform 26.06.

#### Supplementary Table 13 – Immune- and cardiac-exclusive recovery sets and performance

Provided as external files. Sheets from left to right: “Performance” and “Pairwise DeLong”. The “Performance” sheet describes each score by recovery set combination, one row per combination. Variable "recover" names the recovery set: "immune_exclusive" (n = 372) or "cardiac_exclusive" (n = 114). Variable "score" is the prioritization score evaluated, where "OT_genetic_immune" and "OT_genetic_cardiac" are the Open Targets immune and cardiac genetic-association scores, and "our_model" is IGNITE scores. Variable "auc" is the AUC with variables "ci_lo" and "ci_hi" bounding its 95% CI. Variable "ci" is the same interval formatted for display. Variable "n_pos" is the number of positives in that set. The “Pairwise DeLong” sheet describes each pairwise test. Variable "panel" is the recovery set, and variables "A" and "B" report the compared scores with their AUCs in "auc_A" and "auc_B". Variable "delta" is the difference. Variables "z", "p", "p_holm" and "sig" are the DeLong statistic, its raw p-value, that p-value after Holm-Bonferroni correction within the panel, and the significance annotation, respectively. Construction of the exclusive recovery sets is described in Methods. Indication annotations were derived from the Open Targets Platform 26.06.

#### Supplementary Table 14 – STAT4 evidence dossier

Provided as external files. Sheets from left to right: “Evidence axes”, “Open Targets diseases”, “Knockdown neighborhood”, “Neighborhood summary”, and “Score distribution”. The “Evidence axes” sheet describes the percentile of STAT4 on each of four evidence axes. Variable "axis" is the display label, and variable "feature" is the feature-matrix column the percentile is computed from. Variable "percentile" is the percentile taken among genes with a measured value. Variable "evidence" is the evidence block. Variable "order" is the display order. The Th17 axis used "neglog10_adjp_Th17" where the model uses the signed "zscore_Th17". The rest of the features are among the 24 modeled features of IGNITE. The “Open Targets diseases” sheet decomposes the model's immune-GWAS feature and lists the 46 of STAT4's Open Targets disease associations that carry fine-mapped GWAS credible-set evidence. Variable "disease" is the Open Targets label verbatim, and variable "label" is the shortened form the figure draws. Variable "efo" is the Experimental Factor Ontology identifier. Variables "gwas_credible_sets_score" and "overall_assoc_score" are the credible-set datasource score and the overall association score, respectively. Variable "tag" denotes whether the disease falls inside the immune-system-disorder subtree MONDO:0005046. Variable "contributes_to_gwas_score" indicates whether its evidence rolls into the model's feature. Variable "n_credible_sets" describes how many credible sets it contributes. Variable "best_l2g" is the highest locus-to-gene (L2G) score among them. Variable "credible_sets" is the sets themselves, by Open Targets study-locus identifier separated by "; ". The 21 diseases inside the subtree carry 115 credible sets between them. The “Knockdown neighborhood” sheet describes the six nearest approved immune targets by cosine similarity of their knockdown signature to STAT4’s signatures. Variable "gene" is the neighbor, and variable "cosine" measures the similarity. Variables "mechanism" and "pathway" are their manually curated annotation. Variable "approved_drugs" names every approved drug for the target separated by "; ". They are ordered with immune-indicated drugs and approval first. The “Neighborhood summary” sheet reports the variables "statistic" and "value" for the “Knockdown neighborhood” sheet. Statistics include the focal gene, condition, size of the anchor set, comparison distribution, and the distribution's 95th percentile and median. The “Score distribution” sheet reports variables "gene" and "pu_score" for all 19,502 genes, against which the STAT4 score of 0.963 at rank 13 is positioned. These scores are identical to the corresponding column of Supplementary Table 1. The neighborhood analysis is described in Methods. External annotations were derived from the GWAS Catalog and the Open Targets Platform 26.06 and were not used as model features.

#### Supplementary Table 15 – Nearest approved immune-target knockdown neighbor by condition

Provided as external files. Each stimulation condition is present on separate sheets. Sheets from left to right: “Rest”, “8 h stimulated”, and “48 h stimulated”. Columns are identical for these sheets. Genes carrying a significant on-target $CD4^+$ T-cell knockdown and their nearest approved immune-drug target in knockdown-signature space are described in each sheet, one row per perturbed gene, ordered by descending similarity. Variable "gene" is the perturbed gene. Variable "kNN_max_score" is the maximum cosine similarity of its knockdown signature to any gene in the anchor set, which are approved immune targets. Anchor set includes 105 genes at rest, 112 at 8 h and 112 at 48 h. Variable "nearest_immune_target" names the anchor gene achieving the maximum cosine similarity. Variable "is_approved_target" marks the rows where the perturbed gene in “gene” variable belongs to the anchor set, which are scored against the remaining anchors rather than excluded. Variable "drug_name" names approved drugs targeting the nearest anchor gene separated by "; ". Variable "immune_indications" are the immune indications annotated to drugs targeting the nearest anchor gene at any development stage separated with “;”. Variable "n_downstream_z3" counts the measured genes whose expression shifts by more than three standard deviations on knockdown. Variable "culture_condition" repeats the sheet's condition so that the three can be concatenated without loss. Drug and indication annotations were derived from the Open Targets Platform 26.06. These values are a post-hoc interpretive annotation rather than a screening score and are not among the 24 modeled features of Supplementary Table 2.
